# Benchmarking of small and large variants across tandem repeats

**DOI:** 10.1101/2023.10.29.564632

**Authors:** Adam English, Egor Dolzhenko, Helyaneh Ziaei Jam, Sean Mckenzie, Nathan D. Olson, Wouter De Coster, Jonghun Park, Bida Gu, Justin Wagner, Michael A Eberle, Melissa Gymrek, Mark J.P. Chaisson, Justin M. Zook, Fritz J Sedlazeck

**Author notes:** equal contributing (Justin M. Zook, Fritz J Sedlazeck).

## Abstract

Tandem repeats (TRs) are highly polymorphic in the human genome, have thousands of associated molecular traits, and are linked to over 60 disease phenotypes. However, their complexity often excludes them from at-scale studies due to challenges with variant calling, representation, and lack of a genome-wide standard. To promote TR methods development, we create a comprehensive catalog of TR regions and explore its properties across 86 samples. We then curate variants from the GIAB HG002 individual to create a tandem repeat benchmark. We also present a variant comparison method that handles small and large alleles and varying allelic representation. The 8.1% of the genome covered by the TR catalog holds ∼24.9% of variants per individual, including 124,728 small and 17,988 large variants for the GIAB HG002 TR benchmark. We work with the GIAB community to demonstrate the utility of this benchmark across short and long read technologies.

## Introduction

Tandem repeats (TRs) are direct head-to-tail repetitions of a DNA motif^1^. The constituent motifs can be exact copies or contain mutations relative to a consensus motif. A TR region in the genome may contain any number of copies of a single consensus motif, multiple abutting motifs, and even nested repeat structures. TRs are typically classified into subtypes based on the motif’s length (short tandem repeats (STR), 2-6bp^2^; variable number of tandem repeats (VNTRs), 7-100bp^3,4^) or by their genomic context or function (e.g. alpha satellite repeats in centromeres or rDNA repeats). Many TRs are highly polymorphic, with allelic diversity across the length of their expansions/contractions as well as mutations within their motifs or near their boundaries^5,6^. These polymorphisms can be found across populations, as *de novo* changes across generations^7^, and contribute to somatic variability^8^. A subset of TR alleles have been associated with human phenotypes, including many neurological or neurodegenerative diseases^9–12^. The variability of TRs also makes them an important tool in forensic science^13,14^, with the most notable example being TRs characterized by the Combined DNA Index System (CODIS). Together, these properties of TRs highlight the necessity of fully interrogating TR regions of the genome for evolution, medical, and forensic studies.

There exist multiple methods for capturing TR diversity in terms of length and sequence composition. Early methods include low-throughput techniques such as Southern blotting, PCR, and repeat-primed PCR^15,16^. These methods are common for identifying the length of the TR in the individual, but often do not aim to resolve detailed information about smaller mutations like SNVs. High throughput short-read approaches have been introduced to determine variants in TRs with disparate levels of completeness^17–19^. While these methods target a substantial fraction of TRs, they are unable to resolve many locations on the human genome, particularly when the TRs are much longer than the read length. Long-read methods have expanded the detection of variants in longer TRs, including VNTRs, and can be used to assemble multi-Mbp satellite repeats^14,20,21^. In fact, novel phased assembly approaches such as the combination of sequencing technologies are now able to resolve many of these difficult regions^22–24^.

Most TR detection methods target subsets of TRs, while some elucidate signals over a larger catalog of predefined TR. Because of the diverse set of sequencing technologies and bioinformatics tools used to capture TRs, it is difficult to account for the relative strengths and weaknesses of analysis choices made. Without a standardized comparison approach, accurately assessing method performance becomes subject to bespoke validation that may lack comprehensiveness and potentially hinder scientific advancement.

High-quality benchmarks serve to advance the development of novel technologies and variant calling methods^25,26^. The Genome in a Bottle Consortium (GIAB) has produced several important benchmarks for single nucleotide variants (SNV), small insertions and deletions (indels), as well as structural variants (SV)^25,27,28^. These benchmarks have had a substantial impact on the community and are regarded as the gold standard for the evaluation of sequencing technologies and variant calling methods^25^. One of GIAB’s recent benchmarks resolved ∼70% of challenging medically relevant genes, with the remaining genes deemed too complex to resolve due to their repetitiveness (e.g. very long tandem repeats and segmental duplications) or the inability for variant comparison tooling to accurately evaluate their variants^29^. Similarly, TRs’ repetitive nature, complex mutational patterns, and allele representation ambiguities often prohibit their inclusion in genome-wide benchmarks^25^.

Variant comparison tools are fundamentally important to genomics benchmarking. These tools aim to determine the shared variation between a baseline truth set and comparison result (true positives or TP) as well as variants unique to each set (false negatives FN (ie. missed) and positives FP (i.e. additional) variants, respectively). There has so far been a co-evolution of variant comparison tools with the benchmarks to which they’re applied. For small variant benchmarks, the recommended variant comparison methods are designed to specifically handle small variants (SNVs and indels). For example, vcfeval and hap.py perform well when comparing most variants less than 50bp in length, but often fail for larger variants (e.g. SVs) and smaller variants near SVs^30^. Similarly, SV benchmarks are accompanied by variant comparison tools designed for variants that are at least 50bp^30,31^. This separation of variants by size prevents previous variant comparison tools’ applicability to TRs since they contain small variants, large variants, and variants with ‘medium’ (10bp-50bp) length. Another challenge TRs pose to variant comparison tools is that the optimal alignment path through repetitive elements is highly sensitive to alignment parameters and small point mutations^32,33^. This sensitivity can cause homologous alleles to produce variants that are shifted in position, split into multiple variants, or - in extreme cases - give rise to different variant types. Thus, to provide the community with a TR benchmark that assists genomic discovery, a variant comparison method that works across variant sizes and overcomes ambiguous representations is paramount.

Here we describe a genomic benchmark for tandem repeats (excluding homopolymers) across the GIAB HG002 individual’s genome. We begin by cataloging 1,784,803 tandem repeat regions (covering 8.1% of GRCh38) from all TR classes and characterizing the regions’ variants across 86 haplotype-resolved long-read assemblies, followed by the curation of the HG002 TR-specific benchmark. This TR benchmark curation resolves 95% of all cataloged tandem repeats containing 124,728 indels (5-50bp) and 17,988 structural variants (≥50bp) in HG002. In building this TR benchmark we highlight insights into TR regions and discuss their impact. We also present an improved approach to variant comparison by expanding the popular variant comparison tool Truvari^30^. These improvements include the ability to compare small (≥5bp) and large variants simultaneously as well as a variant harmonization technique which overcomes variant representation ambiguities across sequencing technologies and variant callers. Furthermore, to assist the interpretation of benchmarking results over these TR regions, we provide a new variant stratification reporting tool named Laytr. These improvements in benchmark tooling and inclusion of smaller variants and SVs together for the first time in a GIAB benchmark enable the exploration of previously under-investigated TR regions. Lastly, we explore the reliability of our benchmark and the commonality of HG002’s alleles. Here pathogenic alleles are not present, nevertheless, many interesting regions highlight that HG002 is an ideal candidate to represent TR diversity among the human population.

## Results

### Cataloging tandem repeat regions

Resources defining subsets of tandem repeats’ (TRs) locations are plentiful^34–37^. However, each differs from the next due to methodological differences or their focus on specific categories of TRs. To obtain a general tandem repeat catalog across the human genome, we collected genomic intervals from nine sources of TR definitions on GRCh38 autosomes and sex chromosomes (Supplementary Table 1). These sources contained between between 5,765 and 1,737,251 TR intervals, covering 0.01% to 4.55% (0.3 to 129 Mbp) of GRCh38. Supplementary Figure 1 shows an upset plot illustrating each source’s contribution to the merged set of candidate TR regions. Notably, there is high discordance between sources as each generally aimed to capture specific subsets of TRs (e.g. VNTR vs. STR).

Inter-source consolidation was performed by filtering to intervals between 10bp and 50kbp in length and extending each interval’s span by 25bp upstream and downstream before merging intervals with ≥1bp of overlap (see Methods). This procedure resulted in 2,171,789 candidate TR regions spanning 10.5% (308 Mbp) of GRCh38. The sequence spanned by each candidate interval was then analyzed by TandemRepeatsFinder^38^ to capture TR motifs and reference copy numbers. Each interval’s annotations were then processed to create the final simplified TR catalog. This processing excluded homopolymer TR annotations, which are generally short and may be overly subject to sequencing errors^6^, and intervals without TR annotations, which lack identifiable motifs. We also collapsed redundant TR annotations and collected per-region summary metrics such as motif purity (see methods). This produced 1,784,804 TR regions spanning 8.1% (236 Mbp) of GRCh38, with 66.2% of TR regions containing exactly one annotation and 31.5% containing between two to five annotations of TR motifs.

For each TR region in our catalog we recorded additional information in addition to the regions’ coordinates and TR annotations within each region; a full description of the catalog’s information is provided in Supplementary Table 2. The first useful piece of information is a recording of the length of the buffer sequence between the TR region’s start/end and its first/last TR annotation. The extension of input intervals’ span by ±25bp aimed to center TR annotations within a buffer of non-TR sequence in order to allow for alignment ambiguities when capturing variation within a region. However, TandemRepeatsFinder was given the entire region’s sequence and therefore TR annotations may have reached into the buffer. A total of 1,377,360 (77.1%) of TR regions have at least 5bp of upstream and downstream buffer. Another 407,444 (22.8%) of TR regions have less than 5bp of buffer at either end (Supplementary Figure 2).

The information recorded in the catalog also includes an overlap flag which describes the proximity of TR annotations to one another in each TR region (see Methods). This overlap flag estimates 92.0% of TR regions having ‘simpler’ motif patterns including 68.9% of TR regions containing isolated, non-overlapping annotations, and 19.0% having some combination of ‘parent/nested’ annotations. Supplementary Table 3 holds a pivot table summarizing overlap flag counts.

TR Regions with tandemly duplicated interspersed repeats (e.g. short interspersed nuclear elements (SINEs)) can be identified by TandemRepeatsFinder. While this patterns fits the algorithmic definition of tandem repeats in that they are at least two adjacent copies of highly homologous sequence, they may not be subject to the same mechanisms of contraction/expansion (i.e. slip-strand mispairing) seen in a more restrictive definition of tandem repeats^39^. Therefore, information recorded in the catalog also leverages RepeatMasker^40^ results to report the presence of interspersed repeat elements. In total 100,361 (5.6%) TR regions were found to have interspersed repeat elements, which are mostly SINEs (89,671).

Along with these repeat specific annotations, we labeled TR regions that intersect TRs with previously reported importance, namely CODIS^41^, pathogenic TRs^17,35,42–45^, and VNTRs that shape human phenotypes^46^. We found all 53 CODIS TRs were present in the catalog. However, four CODIS TRs were paired into two TR catalog regions: DYS389I / DYS389II, which has an overlapping structure^47^; DYS461 / DYS460^48^, which are reported as being 105bp apart in GRCh38 and spanned by a single ‘TCTA’ motif in the catalog. The TR catalog also includes 67 out of the collected 68 known pathogenic TR regions (see methods). The single pathogenic TR lacking representation in the catalog was VWA1 as it had no overlap with the initial TR intervals. Furthermore, two TR regions in the catalog span multiple known pathogenic TR loci (ARX: 2 loci, HOXA13: 3 loci). Finally, of the 118 VNTRs reported to have a phenotypic impact, 117 overlap the catalog’s TR regions with the single missing locus being ZFP28. However, four of the reported VNTRs spanned multiple TR regions (LPA overlapped 22 TR catalog regions; NEB overlapped 16; 4 for DMBT1; 2 for ZNF181). These loci were therefore not labeled in the TR catalog.

To estimate the accuracy of the motif annotations generated for the TR catalog, we compared the derived motifs to those reported from the aforementioned overlapping pathogenic and phenotypic TR sources. Overall we saw high concordance from the pathogenic TRs with 59 (92.1%) having exact matching motifs to the catalog’s and two more having high similarity but with a more parsimonious motif sequence (see methods). However, only 31 (27.4%) of the phenotypic VNTRs had motifs matching the catalog. The lower concordance of the phenotypic VNTRs was expected given the known difficulties of representing VNTR consensus motifs^6^. Therefore, we also checked how many phenotypic VNTRs’ motif lengths were within 1bp of the catalog’s motif lengths and found 63 TRs (55%) satisfied this condition.

We also annotated our TR catalog’s overlap with genes. This identified 1,109,281 (62.2%) TRs as having overlap with genes, including 831,837 (46.6%) with protein-coding genes. Additionally, we checked for enrichments of TRs relative to promoters and gene features (Supplementary Figure 3). Promoters were observed to overlap TRs significantly more than expected by random chance (permutation test p-value<0.001) with 6,921 of 29,598 promoters (23.4%) intersecting a TR. For protein-coding gene transcripts, the TRs overlap less frequently than expected (p-value <0.001). Intersecting protein coding transcripts’ coding sequences separately from their untranslated regions found both sets’ intersection to TRs to be significantly lower than expected with 25,793 and 38,427 intersections, respectively (p-value <0.001). All exonic regions (N=224,041) rarely overlapped TRs with only 58,966 intersections compared to the permutation test’s mean and standard deviation of 74,610±795. These results are consistent with previous reports on properties of TRs occurrence in relation to genomic features^49,50^.

To visually inspect the TR catalog, we built a self-organizing map (SOM) using kmer-featurization of each TR region’s 4-mers (Figure 1a). This unsupervised machine learning algorithm groups similar sequences by projecting the 256-dimensional space that is 4-mer frequency per region to a two-dimensional space while preserving the topology of TR sequence composition. To assist interpretation of the SOM, we mapped 100 randomly selected TRs which were found to have SINEs, 62 known pathogenic, and 113 phenotypic VNTRs TR regions from the catalog to the SOM (Figure 1a). We first observe that TRs with SINE sequences largely fall into a single neighborhood of neurons (top middle). Additionally, 50 known pathogenic repeats generally fall into three main neighborhoods of the SOM due to having similar motifs (e.g. 25 known pathogenic repeats have CGG/CGC motifs and cluster in the CGS neighborhood). On average 2,588 TR regions map to each neuron in the SOM (Figure 1b, Supplementary Figure 4). This highlights not only the utility of the SOM to separate and visualize TRs by sequence context, but also suggests high TR sequence diversity.

**Figure 1.**
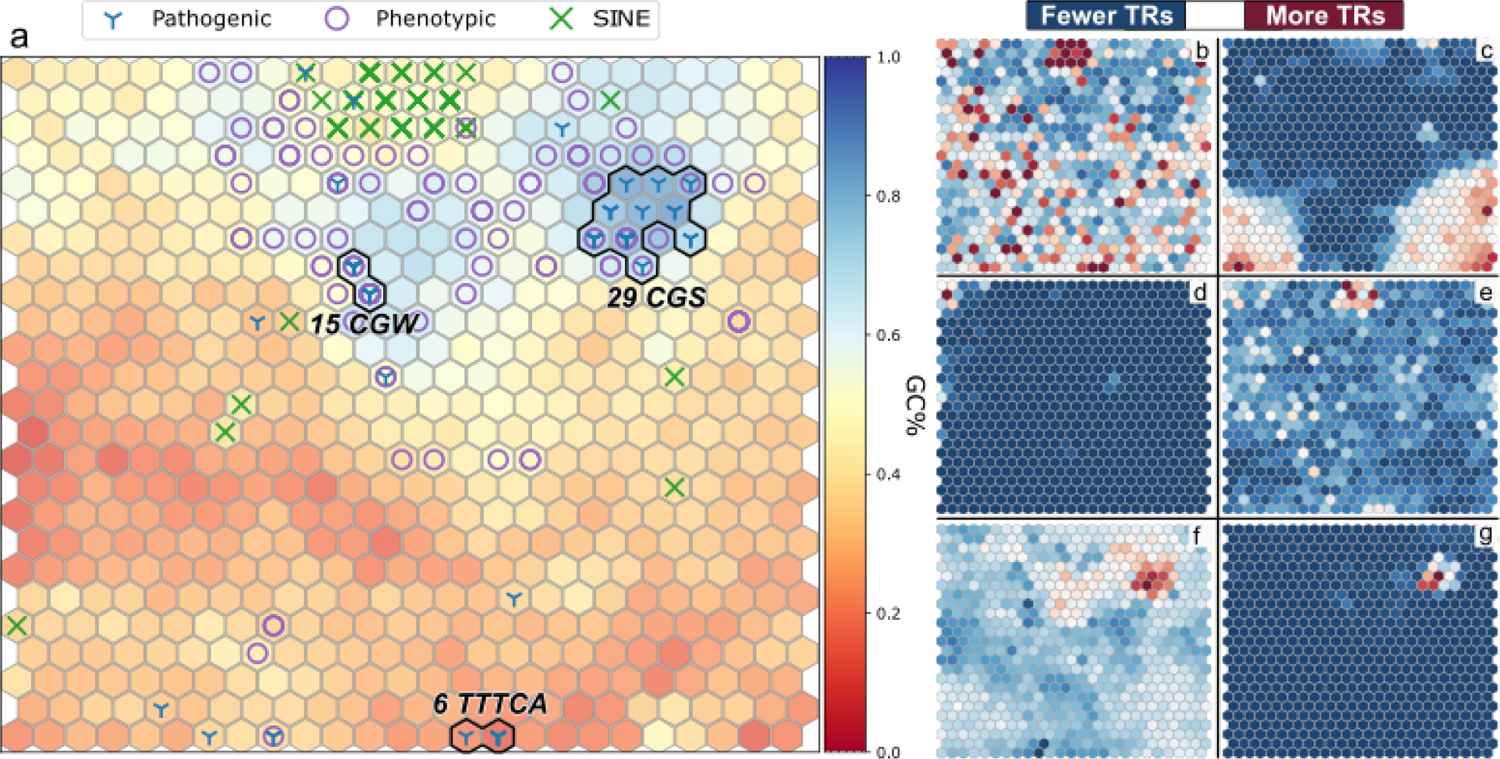
Sequence contexts of tandem repeat catalog. **(a)** Self-organizing map using 4-mer frequencies per-region with hue indicating mean GC percent. Dense known pathogenic neighborhoods are annotated with their most common motif using IUPAC codes (S=G|C, W=A|T) **(b)** Number of TR regions per neuron. **(c)** Average percent of TR regions’ sequence annotated as a homopolymer. **(d)** A neighborhood of microsatellites is observed on the top left. **(e)** Visualisation of segmental duplications shows clustering in similar sequence contexts as SINE elements in (a) at the top-center. **(f)** Map of TRs intersecting genes. **(g)** TRs overlapping promoters.

During the cataloging process regions with only homopolymer annotations were removed. However, sequences with lower complexity may have been annotated as both a homopolymer run and, for example, a dinucleotide repeat. In these cases the homopolymer annotation was removed and dinucleotide annotation was preserved. In order to identify these ultra-low complexity sequences in the catalog we recorded the percent of a region’s sequence that was annotated as a homopolymer run. Figure 1c shows the mean percent of TR regions’ sequence that is annotated as a homopolymer and exposes two dense concentrations in the bottom corners corresponding to neighborhoods of ultra-low complexity sequence.

We further plotted TRs based on their intersection to four reference annotation tracks to highlight their representation in our catalog. Microsatellites (Figure 1d) form a neighborhood in the top left, suggesting they have a distinct sequence context. TRs overlapping segmental duplications (Figure 1e) are mainly concentrated in the SINEs neighborhood. TRs intersecting genes (Figure 1f) and promoters (Figure 1g) are highly concentrated in GC-rich neighborhoods and, unsurprisingly, overlap the bulk of known pathogenic TRs^51^. These neighborhoods of TR sequence contexts overlapping various reference annotation tracks showcase the diversity and comprehensiveness of the TR catalog.

### Polymorphism across tandem repeats

Having defined a catalog of TR regions, we explored the variation within these regions across three independently derived HG002 assemblies as well as 83 haplotype-resolved long-read assemblies of other individuals from multiple projects and populations^52–54^ (Supplementary Table 4). This exploration helps us understand the genetic diversity across each TR region while refining and annotating our benchmark. The assemblies were processed through Minimap2^55^ to produce alignments, variants called with paftools^55^, and a custom pipeline collected per-haplotype coverage of the assemblies (see methods). From these 86 assemblies, we created a unified set of variants as a project-level VCF (pVCF).

On average, haplotypes had a contig N50 of 35.3Mbp. Assemblies produced by HPRC (N=46), which used hifiasm^56^ to assemble PacBio HiFi reads with parental short-reads, were found to be more continuous (mean haplotype contig N50 44.5Mbp) than those produced by other projects (N=39, N50=27.3Mbp). Furthermore, the HPRC HG002 assembly underwent additional gap-filling, scaffolding, and curation to extend its continuity from an N50 of 86.3Mbp to 121.2Mbp^57^. This set of 86 assemblies from 78 individuals produced alignments covering 96.7% of GRCh38 on average. We classified spans of the reference as being confidently covered by a sample if its haplotypes each produce a single alignment over a span (see methods). On average, the HPRC samples’ assemblies confidently cover 96.2% of GRCh38 compared to 92.3% for the other projects (Supplementary Table 5). Therefore, we chose the HPRC HG002 assembly with super-scaffolding to produce the variant representations used by the tandem repeat benchmark.

Samples produced between 5.7M and 10.2M (mean 7.2M) confidently covered variants depending on their population (Supplementary Figure 5). Despite the TR catalog only covering 8.1% of GRCh38, an average of 24.9% of variants per sample occur within TR catalog regions. The observed enrichment of samples’ variation in TR regions increases with variant length with an average of 21.2% of single-nucleotide variants (SNVs), 36.6% of small insertions or deletions (indels) of less than 5 base pairs (bp), 66.6% of variants between 5 and 50 bp, and 72.2% of variants at least 50 bp long occurring within TRs. Merging variants across samples produced a pVCF with a total of 124 million variants. We again see a significant enrichment of variation in TRs (overlap permutation test p<0.001) with 20.7% of pVCF variants inside TR regions and the same relationship between variant length and TR enrichment (Supplementary Table 6). Since variant counts can be sensitive to alignment parameters (i.e. varying allelic representation), we also summed the length of all non-SNP variants in the pVCF and again found 486Mbp (41.0%) of the 1.1Gbp variant bases are within TR regions. These per-sample and pVCF variant metrics illustrate the highly polymorphic nature of TRs and their disproportionate contribution to genetic diversity.

This enrichment of variants is not uniformly distributed across TR regions. Of the 1.78M TR regions in the catalog, 27.2% have no observed variation across the 86 assemblies, 47.4% have only small indels, and 81.0% have no variant ≥5bp in length. The 19.0% of the TR catalog with at least one variant ≥5bp spans 3.0% of GRCh38 and contains 70% of pVCF variants ≥5bp. We attempted to replicate this observation by intersecting variants ≥5bp from dbSNP v153 across chromosome 1 with the TR catalog and found that while 79.1% of TRs have at least one dbSNP variant of any allele frequency (AF), 74.0% lack a variant ≥5bp with AF above 1%. We then partitioned TR regions by if they contained a variant ≥5bp in the pVCF and performed a permutation test on their intersection with protein-coding genes. This experiment showed that the 1,446,782 TR regions without ≥5bp pVCF variants intersect genes as frequently as expected by chance (p=0.45), while the 337,627 with variants ≥5bp have significantly fewer intersections with genes than expected (p<0.001). Thus, we observed a higher frequency of variants in TR regions compared to the rest of the genome, with larger variants being underrepresented in protein-coding genes.

### Formalizing the HG002 tandem repeat benchmark

To create an HG002-specific benchmark, we leverage assemblies of HG002 from three projects. We then subset the TR catalog to regions that are confidently covered (1x per-haplotype) by the scaffolded HPRC HG002 assembly and an ‘alignment replicate’ of that same assembly created using different alignment parameters (see methods). By analyzing two alignments of the same assembly with different parameters we produce different sets of confidently covered TR regions and variant representations. Restricting to TR regions that are consistently covered by both the primary and replicate alignment excludes regions that may be overly subject to alignment ambiguities. Furthermore, a repeat region was excluded if either haplotype has a break in the assembly-to-reference alignment inside a segmental duplication, a tandem repeat longer than 10 kbp, or a satellite repeat. We also excluded all gaps and homopolymers longer than 30 bp, since assembly-based variant calls are less accurate in these regions^27^ (see methods).

This curated set of confidently covered regions captures 1,706,853 TR regions (95.6% of the TR catalog) for inclusion in the benchmark across the autosome and sex chromosomes (Figure 2a). To assess the sequence composition of the benchmark’s TR regions we again used the aforementioned kmer-based SOM, which clusters regions into neurons. On average, 95.9% (standard deviation ±3.8%) of TR regions per neuron are captured in the benchmark (Figure 2b; Supplementary Figure 6). Some neurons (i.e. collections of TR regions by sequence context) are captured below ≤90% due to their TR regions being excluded from the benchmark during assembly curation. This is observed in the few highlighted (red) neighborhoods from Figure 1c & 1e - which represent TRs by homopolymer sequence and intersection with segmental duplications, respectively - overlapping lowlighted (blue) neurons in Figure 2b. The exclusion of long homopolymers from confidently covered regions causes a relative deficiency in the benchmark’s ability to assess TRs in sequence contexts with ultra-low complexity. Nevertheless, 92.8% of TRs in neurons with ≥15% average homopolymer sequence across regions are still included in the benchmark (Supplementary Figure 6). Therefore, TR regions with ultra-low complexity sequences are still represented. Similarly, 127,682 of 364,589 (35.0%) TR regions overlapping segmental duplications remain included in the benchmark.

**Figure 2.**
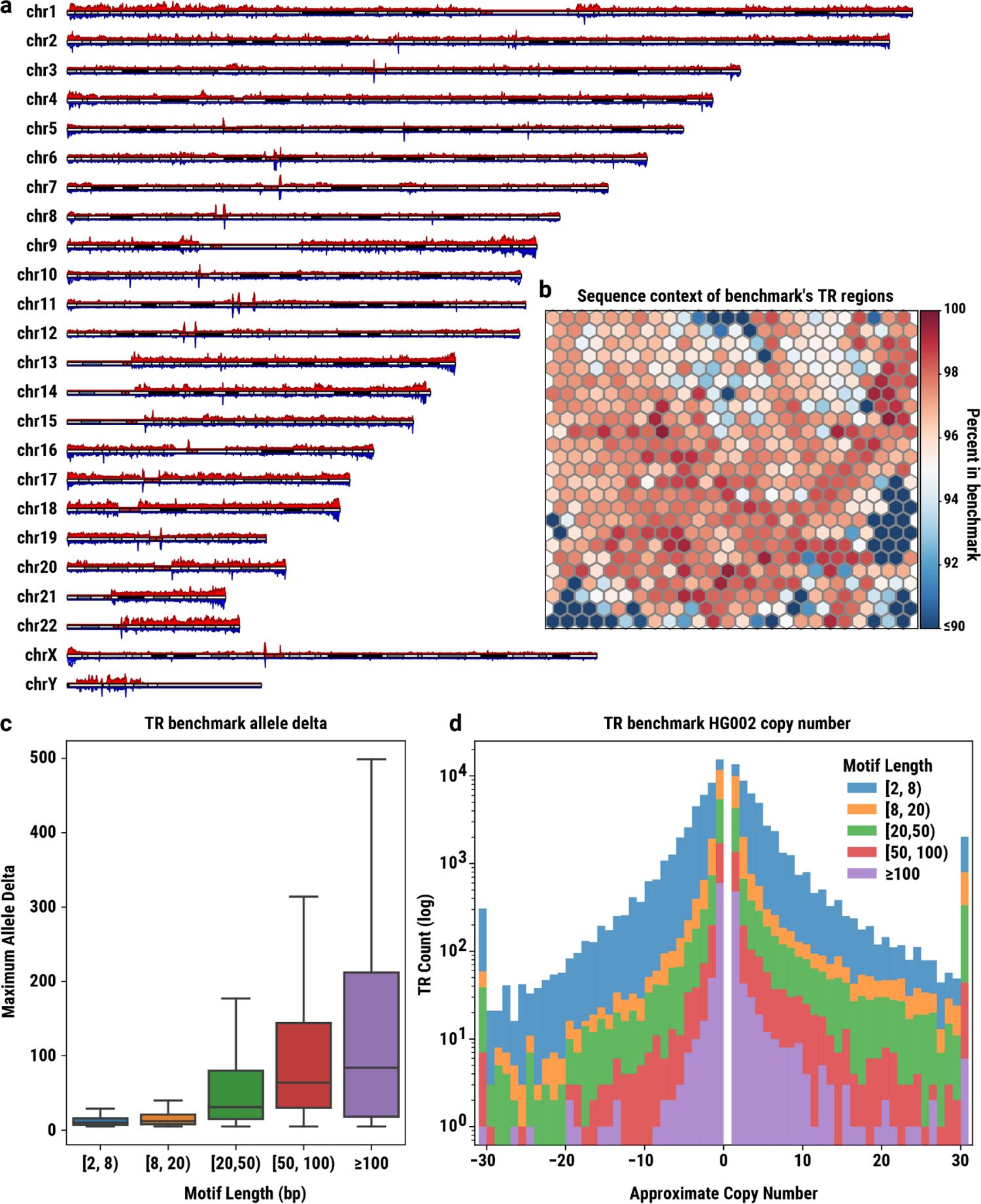
Location, sequence, and length properties of benchmark’s TR regions. **(a)** Karyoplot of TR regions: red (top) TR regions included in the benchmark; blue (bottom) catalog TR regions with HG002 ≥5bp variants. **(b)** SOM Heatmap of percent of TR catalog regions per neuron contained in the benchmark. (**c)** Boxplot of HG002 allele deltas (sum of absolute variant lengths) per region by motif length (lower quartile 25th percentile, upper 75th, center median, extrema 1.5 times interquartile range). At heterozygous regions, the maximum delta is used. **(d)** TR allele delta length per region by motif length. Contractions have a negative delta and expansions a positive delta. Allele deltas greater than 30bp are binned at either end of the histogram.

With the set of HG002 benchmark TR regions selected, we next characterized their variants and confirmed their validity using the HG002 replicate assemblies. We limited our investigation to variants ≥5bp in order to avoid smaller base calling or consensus errors in assemblies. In total, 107,842 (6.3%) of the benchmark regions have ≥5bp variants in HG002, 346,935 (20.3%) have only SNPs and smaller indels, while 1,252,076 (73.3%) are reference homozygous (i.e. have no variants) and can therefore serve as negative controls during benchmarking. We next challenged the confidence of the benchmark’s ≥5bp variants and our ability to compare to these variants using techniques described below, thereby stratifying regions into confident (Tier1) and non-confident (Tier2) sets accordingly. If the alignment replicate of the scaffolded HPRC HG002 assembly and at least one of the two technical replicates of HG002 in the pVCF confirms the presence or absence of the benchmark’s ≥5bp variation, the region is considered Tier1. The remaining regions are considered less confident and labeled Tier2 as they are more likely erroneous due to sequencing errors and/or errors in the assembler’s partitioning of haplotypes. In total, 96.0% (201Mbp span) of the benchmark regions are classified as Tier1, leaving 4.0% (11.7 Mbp) as Tier2. Breaking this down further, a total of 1,616,956 (94.7%, 197Mbp span) benchmark regions have unanimous agreement between the three assemblies (98.7% of Tier1 regions). The main source of TR regions being demoted to Tier2 status (63,671 of Tier2 regions; 93.1%) was due to a failure of both technical replicate assemblies to provide confident coverage over the region, which prevented confirmation of HG002 haplotypes. This includes 845 benchmark regions that had no agreement across the three replicates (1.2% of Tier2 regions). A full table of replicate states’ counts and their assignment to Tiers is found in Supplementary Table 7.

By definition, the set of TR catalog regions included in the benchmark is dependent on the TR regions’ location. However, identifying the exact boundaries of TR sequences is difficult^58^. Our TR catalog is also subject to these difficulties, as illustrated in the above analysis on the distribution of region buffer lengths. To ensure variation included in the benchmark is not overly subject to the regions’ boundaries, we reprocessed the benchmark region selection and tiering processes after expanding all TR catalog regions’ span by 10bp on both ends and compared to the unaltered benchmark. In doing so we find 180 extended benchmark regions were no longer confidently covered by the HG002 assembly, 389 regions were assigned different tiers, and 2,968 had a ≥5bp variant within the 10bp extension. This suggests that only 0.2% of region boundaries in the benchmark could benefit from further refinement and, as a whole, the benchmark regions’ span over variants is stable.

Variant counts and sizes are often dependent on alignment parameters^59^, especially in TR regions. Therefore, it is useful to consider allele deltas, which are the sum of variant lengths per haplotype over a TR region. By comparing the maximum allele delta of HG002 haplotypes per region with the maximum TandemRepeatsFinder annotated motif length collected during catalog creation, we see a slight correlation of longer motifs creating larger allele deltas (Spearman r=0.33, p<0.001, Figure 2c). This pattern can be explained by the commonly accepted stepwise mutation model of TRs, which, among other things, describes expansions and contractions as occurring in whole repeat units (i.e. motifs)^2^. Furthermore, we can approximate the copy number change of TRs in HG002 by dividing the maximum allele delta by the longest annotated motif length (Figure 2d). For example, a 10bp allele delta over a TR region having a 5bp motif comprises a two copy expansion of the TR. We see symmetry across the domain of approximate copy number changes with respect to expansions (positive allele deltas) and contractions (negative allele deltas). However, the range of TR counts shows slightly more expansions (52,658) than contractions (49,014). These analyses illustrate the benchmark captures TR expansion and contractions of multiple copy numbers across motif lengths.

Next, we examined the completeness of the TR benchmark in complex or potentially consequential TR regions by analyzing the properties of TRs that intersect with 5,026 previously reported medically relevant genes (MRG)^7^, known pathogenic TRs, VNTRs with reported phenotypic impacts, and CODIS sites (Table 1). Of the nearly 300k TR regions in the catalog intersecting 4,866 of the MRG, 96.8% are confidently covered by the assembly and included in the benchmark, with 92.8% being Tier1. Interestingly 3,069 MRG have at least one TR benchmark region with HG002 ≥5bp variants which is a subset of the 4,113 having ≥5bp variants in the pVCF (median 7 pVCF variants per-gene). For the 80.6% of pathogenic, 98.2% phenotypic, and 86.3% of CODIS TRs included in the benchmark, 86.0%, 97.2%, and 59.0% are labeled as Tier1, respectively. The pathogenic and CODIS subsets of TRs are generally more difficult to resolve and were classified as Tier2 due to both technical replicates failing to supply confident coverage for confirmation. In total, the technical replicate assemblies were unable to cover all eight Tier2 pathogenic TRs, 17 of the 18 Tier2 CODIS, and 11,240 (95.7%) of the Tier2 TRs intersecting MRG. Nevertheless, 64,386 of 66,465 (96.9%) benchmark regions intersecting pathogenic, CODIS, or MRG and containing HG002 ≥5bp variants remain in Tier1.

**Table 1.** Summary of TRs intersecting interesting genomic regions. Medically Relevant Genes (MRG) is a collection of 5,026 previously reported genes across multiple clinical databases. A total of 4,866 (96.8%) MRG intersect at least one TR. HG002 ≥5bp is the count of TR regions with a variant ≥5bp in the benchmark variants. Other ≥5bp is the count of TR regions with a variant ≥5bp in the pVCF from non-HG002 samples.

| | N | benchmark | Tier1 | Tier2 | HG002 $\geq 5$bp | Other $\geq 5$bp |
| --- | --- | --- | --- | --- | --- | --- |
| **Med. Relevant Genes** | 299,633 | 289,964 | 278,225 | 11,739 | 17,201 | 49,219 |
| **Pathogenic** | 62 | 50 | 43 | 7 | 25 | 42 |
| **Phenotypic** | 113 | 111 | 108 | 3 | 49 | 89 |
| **CODIS** | 51 | 44 | 26 | 18 | 25 | 44 |

### Enabling accurate comparison of tandem repeat alleles

Having constructed the benchmark’s regions and variants, we next explored how to accurately compare to this collection of small and large variants since previous benchmarks separated variants by size. For this purpose, we first evaluated existing recommended methods for small variant comparison (rtg vcfeval) and SV comparison (Truvari bench^30^) beside our newly developed variant comparison technique Truvari refine. To assess the performance of comparison methods we leverage the same HG002 assembly, but with different alignment parameters to obtain noticeably different allele representations for complex variants. This setup demonstrates the difficulties of TR variant comparison while allowing theoretically perfect precision and recall since the TR benchmark regions contain only sites well covered by both alignments of the assembly and all discovered variants are derived from an identical input assembly.

In total, the TR benchmark produces 142,716 variants ≥5bp (66,640 deletions, 76,076 insertions) and the alignment replicate produces 161,733 variants ≥5bp (77,751 deletions, 83,982 insertions). When comparing the replicates with vcfeval or Truvari bench we see an F1 (harmonic mean of precision and recall) of 0.839 and 0.862, respectively. The unmatched variants remaining are caused by highly disparate variant representations and other allele size issues. To improve this we have designed Truvari refine which achieves an F1 of 0.993. Comparison metrics are detailed in (Supplementary Table 8). Analyzing the Truvari refine precision and recall of the benchmark’s alignment replicate by Tier shows 1.00/1.00 for Tier1 regions and 0.89/0.86 for Tier2 regions. The reported performance metrics for vcfeval and Truvari bench vary by variant length for both Tier1 (Figure 3a) and Tier2 (Figure 3b) regions, whereas Truvari refine’s Tier1 performance nearly reaches the theoretically perfect precision and recall and Tier2 performance is less variable and consistently higher than the other approaches. Vcfeval reported performance drops off gradually after 20bp in length and the tool doesn’t compare variants over 1kbp by design. Vcfeval is also limited to requiring that haplotypes match exactly, while Truvari’s core comparison approach can allow inexact matches which is demonstrated by Truvari bench reporting higher performance, particularly after 200bp. Of the 886 FNs and 1,088 FPs reported by Truvari refine, all but 2 FNs and 6 FPs are from Tier2 regions.

**Figure 3.**
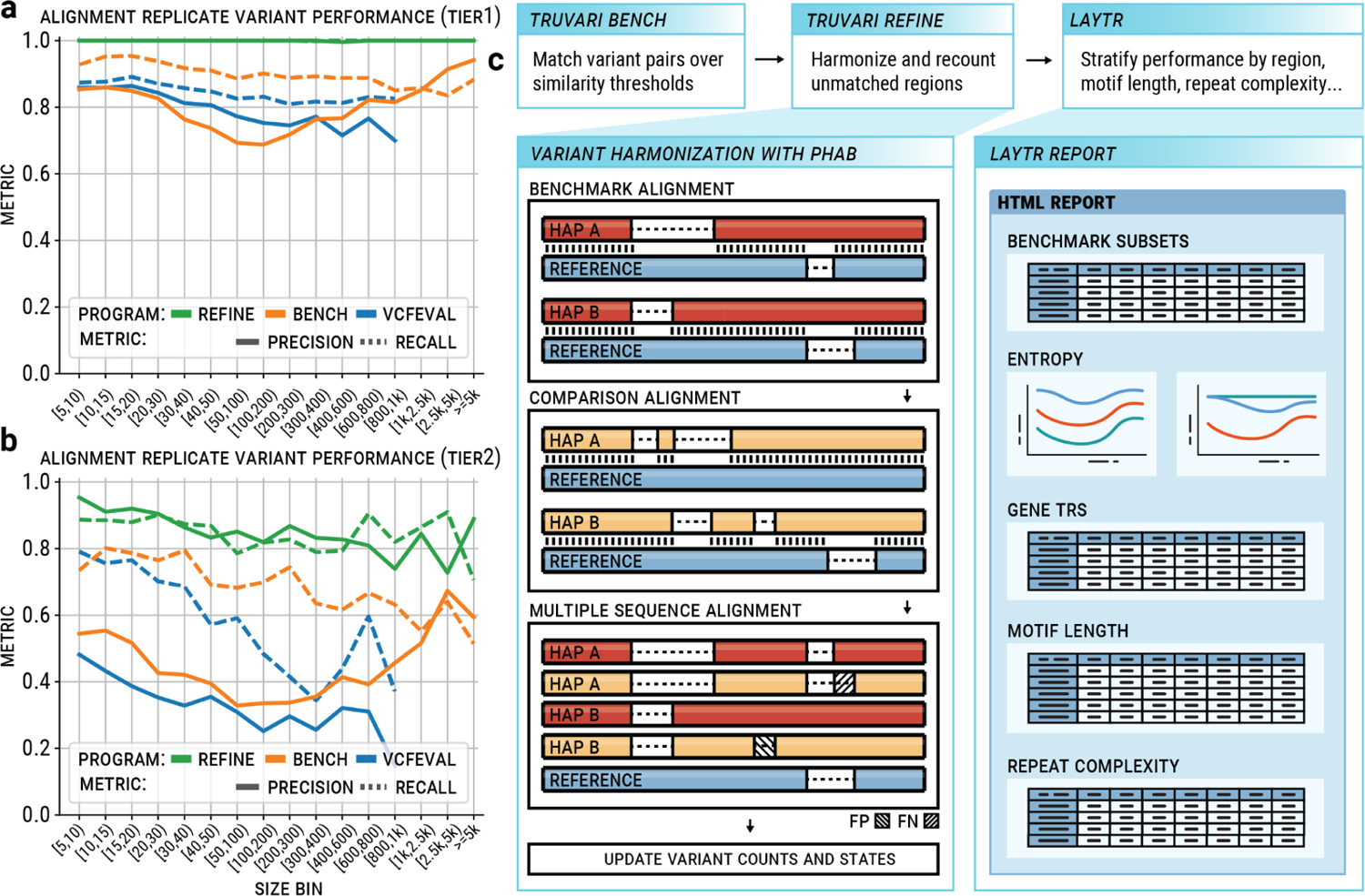
Size regime performance metrics for comparison tools RTG, Truvari bench, and Truvari refine on HG002 TR benchmark against alignment replicate for Tier1. (**a**) and Tier2 (**b**) regions. **(c)** Pipeline schematic of Truvari operations for comparing sequence resolved variants to the TR benchmark. c top - Three commands for creating a benchmarking result and stratification report. c left - Illustration of Truvari phab variant harmonization. c right - Cartoon of Laytr stratification html report.

To accomplish the improvements described above to variant comparison, we expanded the Truvari variant comparison tool’s pipeline. Details of the commands used by the workflow are available in Methods. Figure 3c top shows the principal steps of the comparison procedure. Briefly, to perform variant comparison of TRs, the Truvari bench sub-command’s result is used as input to the new refine sub-command. Refine first identifies benchmark regions with unmatched variants between the baseline and comparison VCFs. Variants in the identified regions are then harmonized with Truvari phab (another new Truvari sub-command) and re-compared with Truvari’s core comparison approach. Summary metrics are then calculated using the original variant counts for un-harmonized regions and the harmonized comparison results for regions which were refined. The per-region summary can then be fed into Laytr - our benchmarking stratification tool - to generate more detailed reporting.

In order to accurately compare TRs for benchmarking, two main challenges needed to be addressed. The first challenge being that the benchmark has variants as short as 5bp which is smaller than Truvari’s previous minimum size of 50bp. The second is that alignment and variant representation ambiguities can be enhanced in TR regions such that 1-to-1 comparison is insufficient. Below we detail the innovations now implemented in Truvari to overcome these challenges as well as report simulation experiments that illustrate the changes’ impact on variant comparison.

Previously, variant comparison with Truvari used a ‘reference context’ approach to calculate sequence similarity in which the alternate alleles of VCF entries being compared are each incorporated into the span of the reference covered by both variants^10^. This produces faux-haplotypes where differences in variant placement (e.g. deleting the first copy vs the last copy of a TR) would have a lesser effect on sequence similarity. While this approach does overcome alignment ambiguities, the similarity scores are inflated by the shared reference context (i.e. reference bases between two variants). This inflation becomes more problematic for smaller variants where reference bases in the faux-haplotypes’ may outnumber the variants’ alternate sequence. Hence, Truvari’s sequence comparison approach was modified to accommodate the possibility of TR expansions and contractions being placed in any position within the reference’s representation of the TR while removing reference bias. This is accomplished by ‘unrolling’ VCF entries’ alternate sequence in a manner akin to variant normalization’s left-alignment^60^. The difference being that left-alignment preserves the order of alternate sequences’ nucleotides whereas unrolling does not. Instead, when unrolling a sequence from its current position to the position of the sequence to which it is being compared, for every base pair the variant is shifted upstream or downstream in another representation, the alternate allele’s 3’ most or 5’ most base pair is moved to the beginning or end of the allele.

To test the accuracy of these sequence comparison techniques, we simulated ∼12 million tandem repeat expansions with between 0% and 30% of bases randomly substituted in the expanded sequence to emulate sequence divergence. We then compared the simulated divergence to the reference context, unroll, and direct comparison sequence similarities (see Methods). On average, unroll sequence similarity differed from the simulated divergence by 0.07 percentage points (p.p.) with ±0.2p.p standard deviation, which is more accurate than reference context similarity which differed by 4.3p.p. ±3.6p.p and direct similarity with 4.3p.p. ±5.5p.p. Supplementary Figure 7 shows how the unroll sequence comparison is more tightly correlated (PearsonR 0.99) to the simulated divergence than reference context (0.67) and direct similarity (0.61).

The second major improvement to Truvari’s comparison is a module for circumventing split variant representations. Split variants are seen when pipelines represent an identical haplotype with a differing number of variants (Figure 3c left). For example, a TR expansion could be represented as a single insertion with two additional copies of a motif or it could be represented as two separate single copy insertions of the motif. While each set of variants describes an identical haplotype, 1-to-1 variant comparisons report deflated similarity metrics as only part of the haplotype’s change in one representation is compared to the full change in the other. To address variant representation differences, a variant harmonization procedure (Truvari phab) was developed which extracts sample haplotypes from VCFs before performing a multiple sequence alignment (MSA) and recalling variants (Figure 3c left).

We simulated 30,000 TR expansions from the TR catalog without interspersed repeat annotations and with an average motif purity of at least 95% (see Methods). Each expansion was then given two variant representations by once inserting the expansion at the beginning of the reference sequence and once inserting the expansion in at least two other positions (see Methods). MSA harmonization reduced the average simulated number of variant positions per-region from 4.11 to 1.87. Since performing an MSA is computationally burdensome, we also implemented an option to perform variant harmonization with the faster wave front aligner^61^ (WFA). While WFA is ∼250× faster than MSA, its independent pairwise realignment of haplotypes produces less parsimonious representations with an average of 1.96 variant positions after harmonization.

Variant harmonization alters their representations and their counts. This becomes a confounding factor for one’s ability to equate performance metrics between runs since the multiple sequence alignment performed causes the final count of baseline variants to depend on the joint alignment created with the comparison’s haplotypes. Therefore, we propose an alternative performance reporting scheme which measures per-region instead of per-variant (Supplementary Table 9). Furthermore, we designed the benchmark to also measure true negatives (ie. regions where no variant ≥5bp should be observed). As we’ve discussed in previous sections, not every TR region has an HG002 expansion or contraction. Therefore, these reference homozygous regions can be leveraged as negative controls in per-region performance metrics. Supplementary Table 10 demonstrates the per-region performance of the three HG002 replicates used for benchmark creation described above. The alignment replicate has a higher balanced accuracy of 0.996 compared to the two technical replicates with 0.990 and 0.989.

In addition to generating accurate comparisons to the TR benchmark, it is also important that comparison results are informative. Throughout the creation of our TR catalog, the HG002 benchmark, and analysis of comparisons to the benchmark we collected many annotations (e.g. presence of interspersed repeats, constituent motifs’ lengths and purity, known pathogenic TRs, intersection to genes, etc). To assist users in leveraging these annotations into informative stratifications, we also provide a new tool named Laytr. Laytr generates per-region performance metrics on 10 standard stratifications of the benchmark (see Methods; Figure 3c right). Laytr joins the Truvari refine region report to the catalog and benchmark data as well as processing the reported regions through the SOM. Laytr produces an html summary for visualization as well as machine readable files. As examples, Laytr reports from the long-read TR caller TRGT^20^ and long-read WGS SV caller Sniffles^62^ are provided in Supplementary Material 1 & 2. Comparing these two reports shows the added benefit of TR specific callers as TRGT has a balanced accuracy across Tier1 regions of 0.98 compared to Sniffles’ 0.53. A considerable proportion of this difference is attributable to Sniffles only reporting SVs ≥50bp whereas TRGT genotypes across all allele lengths. This difference is detailed in the allele delta of the Laytr report which stratifies performance by each region’s sum of variant lengths. This report shows TRGT’s true positive rate (TPR) is consistently at ∼0.98 across all size bins whereas Sniffles only begins to find true positives from 50bp+ at an approximately 0.62 TPR. Complete descriptions of Laytr stratifications and their interpretation are provided in Methods.

The Truvari refine and Laytr methods presented here enable precise variant comparisons especially in TR regions but their concepts can also be applied across other regions of the genome. Furthermore they enable stratification and thus new insights into the performance of different technologies and methodologies to infer variants.

### Insights into the utility of the tandem repeat benchmark

To evaluate the benchmark’s utility in exploring general patterns of TR discovery, we collected variant calls from seven tools that use different sequencing technologies and differing methodologies (Supplementary Table 11) and compared them to our benchmark. These analyses are not definitive assessments of any particular pipeline’s TR discovery capabilities, but instead aim to ensure the benchmark is capable of assisting pipeline evaluations. Four of the tools are TR specific callers, which leverage a TR catalog to discover alleles. Two of the TR callers use long-read sequencing and two use short-reads. The remaining three tools are designed for general whole-genome variant calling (WGS) of different size regimes and discover TR alleles simply due to their previously described prevalence.

As noted above, the number of variants reported after refinement is dependent on the comparison VCF. We find that after harmonization the benchmark’s 139,372 HG002 ≥5bp variants become an average of 150,508 variants across the seven comparisons with a standard deviation of 5,418. However, using Truvari refine’s per-region summaries, we see an average of 106,046 baseline positive (having a variant) TR regions with a tighter standard deviation of 1,191. The remaining variance in region counts is attributable to the MSA ‘resizing’ variants above/below the 5bp minimum length threshold or shifting variants into or out of the TR benchmark’s regions. Still, the tighter standard deviation of per-region summaries allows more consistent comparability between pipelines.

We next searched for patterns in the properties of TRs discovered across these tools stratified by the underlying read length. To do this, we first subset the benchmark to Tier1 regions with HG002 ≥5bp variants that were analyzed across all tools (N=60,325). We next marked TR regions where both short-read TR callers and independently where both long-read TR callers agreed in their benchmarking state (e.g. true positive (TP), false positive (FP), etc). By subsetting to TR regions analyzed by all TR callers and where two tools using the same category of read-length agree, we’re ostensibly limiting the impact of methods on performance in order to query for read-length effects. When comparing the states of regions with short-read caller agreement and long-read caller agreement (N=40,546), we find that 27,807 (68.5%) regions are resolved as TP by both pairs of tools, 12,734 (31.4%) are resolved as TP by only the long-read tools and 5 are resolved as TP by only the short-read tools, though this likely will change as new tools are developed.

When searching for properties of TRs that could indicate if they’re resolved by only the long-read tools (long-read set; LRS) or by all TR callers irrespective of read length (all-read set; ARS), we find four annotations provided by the benchmark with a significant difference (Wilcoxon rank-sum P<0.001). First, the average and standard deviation maximum allele delta (total variant length) of a TP TR region in LRS is 124bp ±471bp compared to 11bp ±14bp in ARS. The TR region’s motif length is 43bp ±74bp for LRS compared to 6bp ±26bp for ARS. The average LRS region span is 593bp ±689bp compared to ARS 125bp ±150bp. And we also find a lower LRS average motif purity (0.91 ±2.5 LRS, 0.95 ±2.4 ARS). In summary, longer changes/motifs over larger TR regions as well as sequence contexts with lower motif purity are resolved more frequently by this set of long-read TR callers than the example short-read TR callers.

In addition to the four TR callers we leveraged three WGS callers and compared the two sets’ discovered TRs. Of the benchmark’s 101,704 Tier1 HG002 ≥5bp regions, the TR callers had at least one TP state in 101,1080 (99.4%) regions and 90,996 (89.4%) regions had a TP state from any WGS caller. A total of 90,812 (89.2%) regions are TP in the sets’ union, 10,296 (10.1%) are found to be TP exclusively by the TR callers, and only 184 are exclusive to the WGS callers. Interestingly, both the long-read and short-read TR callers contribute to the set of regions found exclusively by TR callers at 99.8% and 3.4% of regions, respectively. The increase in TRs resolved by TR callers highlights not only the value of focused TR discovery algorithms but also opportunities for WGS callers to extend the comprehensiveness of their reporting.

As noted above, these results likely will change as new callers are developed and optimized with this benchmark. In addition, the design of the current benchmark and tools does not permit fine-grained analysis of base-level sequence accuracy, which may be higher for short reads for tandem repeats shorter than their read length. Therefore, this analysis should be interpreted as an example of the utility of the benchmark to evaluate several methods rather than an evaluation of how methods using short and long reads differ going forward.

### Diving into the benchmark and its advancements

With the creation of the benchmark complete, the final set of analysis we present aims to answer more general questions about the benchmark. Specifically: how common are the HG002 TR alleles in this benchmark?; What regions and variants are newly contained in this benchmark?; How reliable is this benchmark at identifying errors?

Collecting alleles from multiple genetically diverse individuals allows us to ensure that the HG002 alleles captured in the benchmark are representative of those found across individuals. When checking the 101,704 Tier1 benchmark regions with at least one ≥5bp HG002 variant we find 99.7% have ≥5bp variants present in another sample. However, 7.3% of these regions have non-reference HG002 haplotypes with less than 99% sequence similarity to all other samples’ haplotypes and are therefore considered unique to HG002. Additionally, 181,006 (11%) Tier1 regions lack a ≥5bp variant in HG002 but have ≥5bp variants in another sample. The commonality of HG002 TR variation relative to other samples suggests that variant callers’ performance in discovering HG002 TRs may translate to comparable performance on non-HG002 samples, though it would be valuable to have TR benchmarks for diverse individuals in the future.

While this benchmark contains a comprehensive selection of tandem repeats with respect to length and sequence composition, it’s important to acknowledge the diversity of tandem repeats that may arise at any given locus across individuals (Figure 4). For example, when plotting the allele delta of four CODIS sites across 156 haplotypes from the 78 unique samples in the pVCF (Figure 4a) we observe contractions up to 60bp in length and expansion as large as 100bp. Focusing on just the CODIS PentaE site (Figure 4c) and plotting the unique by length haplotypes with TRviz^63^, we see HG002 allele frequencies (AF) of 0.09 and 0.02 for the paternal and maternal alleles, respectively. These HG002 alleles (10 and 18 copies of the 5bp repeat) are less common than the 8 copy allele with AF of 0.13. The distinction between resolving a locus for a single individual versus all possible allele lengths is especially pertinent to pathogenic repeats (Figure 4b). HG002 represents a healthy individual and thus repeat expansions which are implicated in disease phenotypes are not represented. In Figure 4d we see a wide range of copy numbers across the JPH3^64^ locus but all samples are below the pathogenic length of ≥41 copies. However, in Figure 4e we observe two haplotypes above the ≥50 pathogenic length for TCF4^65^. These TCF4 expansions are up to ∼90 copies of the trinucleotide repeat whereas HG002’s longer paternal allele is only ∼30 copies. Therefore, researchers with an interest in TCF4 would need to account for a pipeline’s ability to resolve the locus in the benchmark as well as the pipeline’s ability to resolve other HG002 loci with longer expansions.

**Figure 4.**
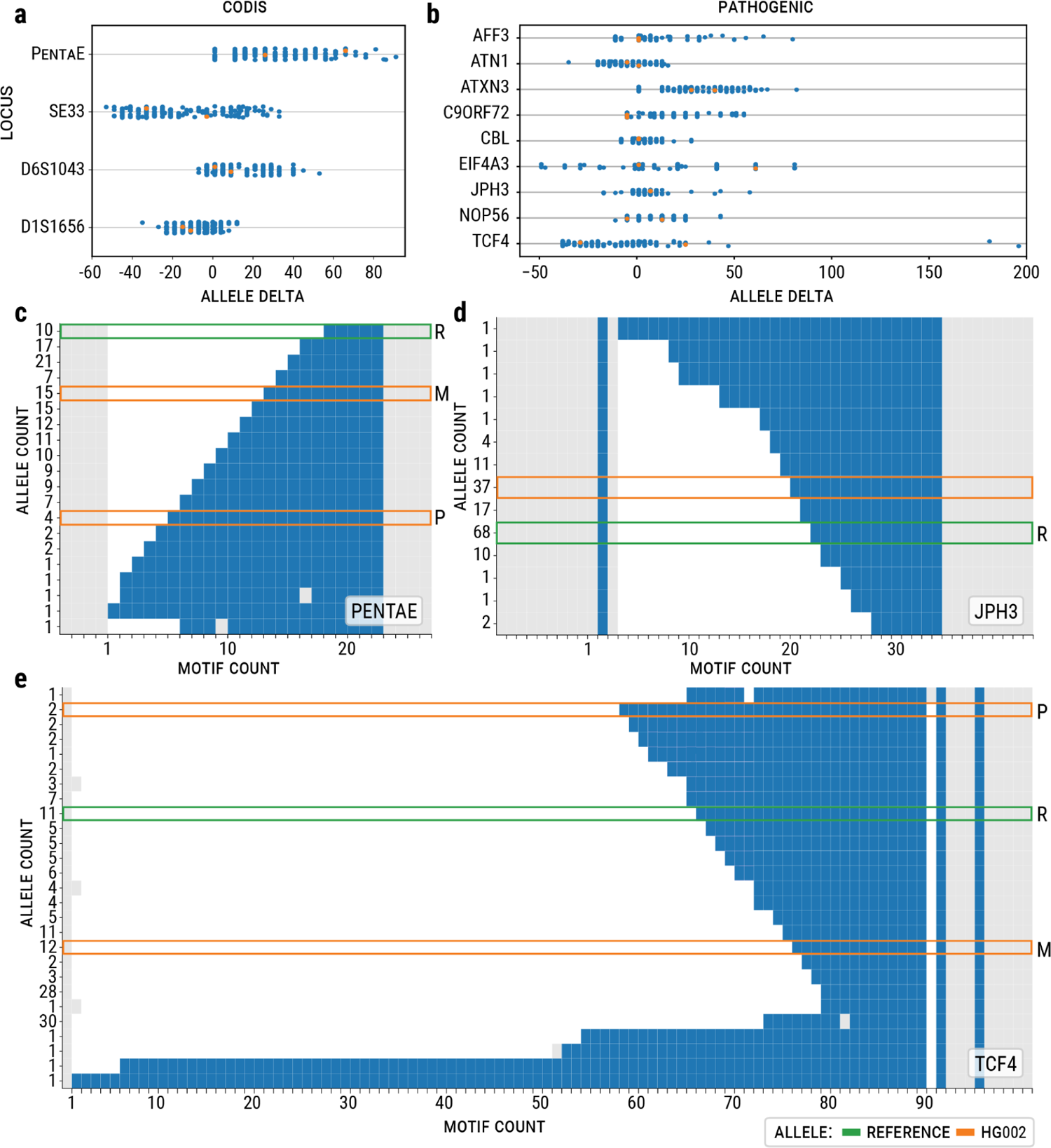
Diversity of tandem repeats over 156 haplotypes at CODIS and known pathogenic loci. (**a**) Allele delta (sum of variant lengths) across four CODIS loci. HG002 maternal and paternal alleles are indicated by orange data points. (**b**) Allele delta of 9 known pathogenic repeats. (**c-e**) distribution of haplotypes over (**c**) CODIS PentaE locus and (**d**) known pathogenic JPH3 and (**e**) TCF4 loci. For **c-e**, each row represents a distinct-by-length haplotype (allele) and the y-label the number of haplotypes with the allele. Blue squares indicate the tandem repeat motifs, gray squares indicate non-motif sequences, and white squares are gaps introduced by multiple sequence alignment. Gray squares upstream and downstream of the tandem repeat are the benchmark’s TR regions buffer sequences. Orange boxes indicate HG002 maternal (M) and paternal (P) haplotypes (homozygous have one box) and green boxes the GRCh38 reference allele (R). Allele count is determined by deduplicating haplotypes by length.

Given that this TR benchmark is preceded by multiple GIAB benchmarks, we next investigated the intersection of the TR benchmark with previous GIAB benchmarks to understand what variants and segments of the reference are newly covered (Supplementary Table 12). Both the GIAB v0.6 SV benchmark^28^ (lifted over from GRCh37 to GRCh38) and the GIAB v4.2.1 small variant benchmark^27^ show large overlaps with the TR benchmark of 95.1% and 84.5% respectively. However, they are constrained in the variants that they report in these regions. This is highlighted by the fact that the TR benchmark includes 126,909 small and 18,720 large variants, which is much higher than previous benchmarks (e.g. v4.2.1 small variant benchmark: 89,029 small and 204 larger variants). Thus, the benchmark’s Tier1 regions include novel variants that are otherwise not captured in existing benchmarks and present opportunities for the community to improve variant calling methods.

To further gain insights into the validity of these variants we collected seven variant callsets across multiple sequencing technologies and analysis techniques (SNV, SV and TR calling methods) (see Supplementary Table 11). We investigated if TR regions in the benchmark where Truvari refine did not assign a TP or TN state to any caller’s result could be a signal that the benchmark’s variants may be incorrect. In total we find 875 (0.05%) TR regions in which all callers disagree with the benchmark. Of these benchmark regions, 866 are also found to be contradicted by the comparison to the high quality GIAB v4.2.1 small variant benchmark^27^. These regions are therefore unlikely to hold accurate descriptions of HG002 TR variants and should be deprioritized for benchmarking. Fortunately, 473 (54.0%) of these suspect regions were already demoted to Tier2 status by the quality control techniques employed during creation of the benchmark and the 402 remaining regions were demoted to Tier2 status in the final published benchmark.

To further ensure the reliability of the benchmark’s ability to identify errors, we manually curated a random selection of Tier1 TR regions with common FP and FN results from callers. We sampled 20 of the 866 (0.05%) Tier1 TR regions where only one comparison callset matched the benchmark. We found that eight appeared clearly correct in the benchmark, and the comparison callsets were inaccurate due to long homopolymers or mapping issues due to a gene conversion-like event and a false duplication in GRCh38. Another four were in TR regions that spanned longer than the length of short-reads and where long-reads were noisy, however the benchmark appeared correct or within one tandem repeat unit of the correct size, but the exact length of the variant was unclear due to noise in the long reads. For another three, the benchmark was off by one or two bp in homopolymers or dinucleotide repeats. The remaining five were incorrect in the benchmark due to the super-scaffolded HPRC assembly having collapsed haplotypes caused by the assembler failing to separate the heterozygous events in an otherwise highly homozygous region, as has been observed previously^27^. Because the benchmark was correct or close to correct even for these variants where most callers disagreed, we kept this as the v1.0 version of the benchmark without further refinement

## Discussion

In this paper we focused on establishing a novel tandem repeat (TR) benchmark to promote the detection and inclusion of variants across tandem repeats in research and clinical settings. This was accomplished by consolidating multiple tandem repeat annotation sources and producing a TR catalog of 1.7 million regions spanning the human genome. Based on 86 haplotype-resolved assemblies, including three HG002 assembly replicates, we were able to form a variant catalog of events ≥5bp in length, which branches across the traditional variant size-regime categories (indel vs. SV). The importance of this benchmark is highlighted by the fact that despite only ∼8.1% of GRCh38 being covered by the TR catalog, its regions include a disproportionate number of variants (24.9% of variants per-sample). In fact, 66.6% of variants between 5 and 50bp (i.e. indels) and 72.2% of variants of 50bp+ (i.e. SV) occur within tandem repeats, implicating an outsized contribution of TRs to genetic diversity and the importance of comprehensive TR characterization for genomic studies.

Many TRs were excluded in previous GIAB benchmarks due to the technical problems associated with both characterization and benchmarking caused by tandem repeats^25^. However, in this study we’ve addressed many of these prohibitive issues and included new spans of the reference in the benchmark. Furthermore, by unifying the previously segregated small and large variants, new alleles are present in the TR benchmark. This novel TR benchmark across allele sizes and sequence contexts demanded a novel approach for variant comparison to ensure accurate reporting of performance metrics. This required the extension of Truvari variant comparison capabilities including an improved sequence comparison technique and variant harmonization procedure to reduce representational ambiguities between callsets. These advances in comparison methodology also required a novel approach to collecting performance metrics in the aforementioned per-region counting metrics. While counting by variants is still useful, assigning states to entire regions was shown to allow more consistent comparison between benchmarking results.

Another unique aspect of this benchmark is the introduction of true negative regions, which are tandem repeats where HG002 contains a reference allele. This is particularly interesting because the complexity of tandem repeats can lead to sequencing errors, misalignments or other sources of noise which variant identification methods need to handle. The simultaneous assessment of true positive, true negative, false positive and false negative thus forms a comprehensive assessment of variant identification methods across the next frontier of genomics and likely clinical genomics as more diseases are linked to tandem repeat mutations.

To extend the utility of comparison to this benchmark, we further provide multiple annotations over the TR catalog and benchmark regions as well as the new tool Laytr to facilitate stratification of benchmarking results. The utility of these annotations were primarily demonstrated in the analysis of multiple tandem repeat and other variant discovery methodologies in which we highlight the contribution of specialized tandem repeat callers. Furthermore, we reported general patterns of TR callers’ contribution when averaged within the read-length they leverage. By subsetting to regions analyzed by all tools and with agreement across callers we limited the analysis to a very conservative set of regions. Fully investigating a pipeline’s capabilities would require a more fine-grained analysis that is best directed by their maintainers and our benchmark hopes to assist those efforts. Therefore, we caution over-interpreting these results as they aren’t representative of any particular pipeline’s limitations. Instead, our aim in these experiments is only to exemplify the utility of the benchmark in hopes that developers of TR discovery pipelines may fully explore the relative strengths and weaknesses of their technologies.

We incorporated into our TR catalog information on known pathogenic and phenotypic tandem repeats together with CODIS tandem repeats that are often utilized for forensic identification. Thus, the utility of this benchmark is not only for bioinformatic developments and human genomic sequencing centers but also for other branches of science such as forensic and evolutionary studies. Still, some of the most complex TR were excluded due to lack of confirmation from the replicate assemblies and were classified as Tier2. This includes e.g. 18 Tier2 CODIS regions across autosome and sex chromosomes. Tier2 regions are considered less reliable for comparison and calculation of performance metrics in this benchmark but still likely harbor interesting variation that should be further investigated and improved upon.

As the HG002 cell line is only a single individual, we investigated the allelic diversity across multiple samples to ensure tandem repeats in the TR benchmark are typical of that expected across a diverse set of individuals. Overall, 99.7% of the regions that carry a mutation in HG002 also carry a non-reference allele across the other 83 assemblies assessed. However, it is important to note that HG002 represents a healthy individual. Thus repeat expansions which are implicated to disease phenotypes are not represented. However, the benchmark’s wide range of allele lengths across sequence contexts (Figure 2) should still serve as a useful test of variant callers for researchers investigating specific loci. As shown in Figure 4, HG002 represents the human population diversity well across different pathogenic variants. Because TRs are so diverse, we provide the pVCF containing 86 samples with 78 unique individuals for researchers to compare against. While only HG002 has been curated and serves as the full benchmark, the other samples may be of descent assistance as controls.

Two challenges not tackled in this benchmark - and thus are a future aim - is the inclusion of ‘novel’ TRs and mosaic variations. First, our work here focuses on TR regions defined by the reference, but TRs also form at loci not initially represented in the reference genome^66^. This could be the source of future work to explore these for benchmarking the ability of methods to identify novel TR regions. Second, mosaic variants often occur at low frequency within a sample and are likely enriched in tandem repeat regions as they might form the basis of the observed diversity^67^. Given our focus on the reliability of our Tier1 regions we utilized assemblies to compile the benchmark’s variants. Thus, low frequency mutations are not reported as they are not produced by diploid assemblies. Characterizing mosaic variants in the individual or cell line will require a deeper interrogation of error modalities of each technology in TRs since it can be hard to distinguish sequencing errors from mosaic variants and errors in repeats can be correlated across technologies. Nevertheless, this is an important future area of work.

Overall this benchmark addresses challenges that have faced the genomic community and promotes the exploration of tandem repeats. The novel methodologies developed here for comparison of variants across tandem repeats will push genomic and genetic research forward by closing a long open gap in existing benchmarks. The benchmark further promotes the notion of identifying all variant sizes simultaneously since small and large variants are present and can now be properly evaluated. We hope this work provides significant assistance to studies’ ability to generate comprehensive whole genome analysis that spans variant sizes and locations.

## Methods

All custom scripts described below are available in the github repository https://github.com/ACEnglish/adotto/. Parameters for software used in analysis are presented in Supplementary Table 13.

### Definition of tandem repeat catalog

Tandem repeat bed files from nine sources were collected^6,17,20,36,51,68,69^ (Supplementary Table 1). Intervals were filtered to exclude spans below 10bp or over 50kbp before performing intra-source merging and boundary expansion of ±25bp using bedtools^70^ v2.31. Inter-source merging was then performed, again using bedtools. A custom script then pulled reference sequences spanned by intervals and processed them with TandemRepeatsFinder v4.09.1^38^ and RepeatMasker v4.1.4^40^. RepeatMasker hits with scores of at least 225 were recorded. TandemRepeatsFinder annotations were grouped per-region and had redundancies removed using a custom simplification script. The simplification script removes annotations of homopolymer motifs and annotations which overlap a longer spanning annotation unless one of the following conditions is met: the boundaries of the shorter annotation are entirely within the span of a single motif from the longer spanning annotation; the shorter annotation boundaries are directly adjacent to a single motif copy of the longer spanning annotation. These two conditions preserved annotations with a parent/nested repeat structure and annotations with unclear start/ends between plausibly adjacent repeats, respectively. TR regions without TandemRepeatsFinder annotations after simplification were removed. Information about each region’s number of annotations, the overlapping relationship of annotations, percent of span annotated, RepeatMasker highest-scoring hit class, intersection with Ensembl v105^68^, intersection with CODIS^41^ and known pathogenic repeats were recorded before uploading the catalog to Zenodo^71^. Full descriptions of the catalog’s columns and definitions can be found in Supplementary Table 2.

### Interval permutation tests

regioneR^72^, an R package for permutation tests on genomic regions, was reimplemented in rust (https://github.com/ACEnglish/regioners). The software counts the number of intervals in a bed file - A which overlap an interval in bed file - B and then performs --num-times permutations where - A intervals are shuffled using a circular randomization strategy before recounting the number of overlaps. The circular randomization strategy shifts all intervals per-chromosome downstream by a random number of bases. If an interval’s new boundaries fall beyond the chromosome’s end, they are ‘circularized’ and wrapped back to the chromosome’s start before continuing the downstream shift. The mean and standard deviation of permutations’ overlaps are then calculated and compared to the initially observed overlap counts to compute a p-value.

Runtimes were collected for ∼29k regions intersected against ∼1.7m on 4 cores on a MacBook Pro. Running 100 permutations on regioneR took ∼1,292 seconds. Running 1,000 permutations of regioners took 3.8 seconds. Therefore, performance benchmarking between regioneR and regioners showed an ∼4,000x improvement in runtime.

The promoter bed file was generated from the UCSC table browser EPDnew track on GRCh38^36,73^. We downloaded gencode v35^74^ and extracted features with protein coding biotypes. Subsets per-feature were collected from the gencode gtf, converted to bed format, and consolidated using bedtools merge. GRCh38 autosomes, chrX, and chrY were analyzed after excluding gap regions as defined by UCSC table browser^36^ via masking by regioners.

### Collection of and motif comparison to known pathogenic repeats and phenotypic VNTRs

Known pathogenic TRs were collected from STRipy^35^, gnomAD^42^, Stevanoski^43^, Dolzhenko^17^, Pellern^44^, Tan^45^. TRs were then consolidated by gene name and overlap of their genomic coordinates was manually ensured. Genomic coordinates per TR were then selected by preference: STRipy, gnomAD, Stevanoski, Dolzhenko, Pellern, Tan. This consolidation resulted in 68 total loci. Additionally, 118 VNTR length protein-coding repeat polymorphisms were collected from Mukamel et.al^46^. The final consolidated set of 186 tandem repeat loci is available in Supplementary Table 14. To compare motifs between the reported known pathogenic and phenotypic repeats and the TR catalog, sets of motifs’ forward and reverse complement sequences were generated and intersected. If any motifs were present in the intersection, the site was considered matched and as such accurately annotated. Furthermore, we allowed ‘Ns’ in the reported motifs to match to any base. Finally, we performed manual inspection of the known pathogenic repeats and found, 4 additional TR regions in the catalog did not match with the above criteria due to a non-parsimonious representation of the reported motifs e.g. TMEM185A’s reported CCG motif is equivalent to the catalog’s CCGCCG and GCN equivalent to GCAGCGGCG for ZIC2.

### Self-Organizing Map

The reference sequence spanned by each region in the TR catalog was put through kmer-featurization using k=4. To generate the kmer-featurization of a sequence, an array of size 4**k is created to hold the observation count of each possible kmer of length k given 4 nucleotides. A kmer is indexed to its array position by taking the summation of NUCS[kmer[pos]] << pos * 2 for pos in len(kmer). Where ‘NUCS’ is a dictionary of {’A’:0, ‘G’:1, ‘C’:2, ‘T’:3}, ‘<<’ is a left bitshift operation, and ‘pos’ is the 0-based index of each nucleotide in the kmer. This kmer-indexing is repeated for each kmer in a sequence being processed and repeated over the sequence’s reverse complement. The array of kmer counts is then normalized by the number of kmers analyzed ((len(seq) - k + 1) * 2) to create a kmer-frequency array. The TR region sequence kmer-featurization generates a NxM matrix where N is the number of regions processed and M is the length of the kmer-frequency array. This matrix is then processed by MiniSom to train a self-organizing map with a T25×25 network of neurons having hexagonal topology. See Supplementary Table 13 for hyperparameters. MiniSom is available from (https://github.com/JustGlowing/minisom).

### pVCF generation

Haplotype-resolved long-read assemblies from 3 sources^52–54^ were collected (Supplementary Table 4). The HPRC HG002 assembly which was further processed to increase continuity^57^ was used in place of the original HPRC HG002 assembly. Each haplotype was aligned using minimap v2.24^55^ and variants called using paftools, which is packaged with minimap. Custom scripts were used to generate per-haplotype coverage bed files. BCFtools v1.12^75^ along with custom scripts were used to perform intra-sample merging, inter-sample merging, and finally allele depth information added to the pVCF using the aforementioned coverage bed files. Coverage bed files per-haplotype per-sample were intersected to identify confidently covered regions of the genome covered by exactly one alignment per-haplotype for the autosome. chrX confident regions were also defined as one alignment per-haplotype for female samples as well as for pseudoautosomal regions of chrX for male samples. chrY confident regions were not collected for female samples and non-pseudoautosomal regions of chrX for male samples were checked for one alignment across haplotypes. The 86 sample pVCF called against GRCh38 is available on Zenodo^76^. The GRCh38 version used did not include alternate loci or decoy sequences (https://ftp-trace.ncbi.nlm.nih.gov/ReferenceSamples/giab/release/references/GRCh38/GCA_000001405.15_GRCh38_no_alt_analysis_set.fasta.gz). Enrichment of variation was tested with regioners (described above) using the pVCF reformatted to bed format and the TR catalog v1.1 with shuffle randomization. Intersection of TR regions with protein coding genes was performed by separating TR regions with at least one variant ≥5bp from TR regions without any variations. Each set was then intersected to Ensembl v105^68^ protein coding gene transcripts with regioners using per-chromosome circular randomization.

### Definition of benchmark regions and variants

The HPRC HG002 alignment coverage bed files built as described above in the pVCF generation step were subset to spans of the reference covered by a single contig per-haplotype using custom scripts. This script only includes regions singly covered by each haplotype (i.e. diploid coverage) for autosomes and 1x coverage for sex chromosomes with the exception of the chrX pseudoautosomal regions as provided by dipcall^77^ which were subset to spans covered singly by each haplotype. Furthermore, a second alignment of the HPRC HG002 assembly was performed using dipcall and custom minimap2 alignment parameters designed to align across complex variants and larger SVs, originally used for the MHC region^78^ (Supplementary Table 13). This produced a second VCF as well as a bed file of regions covered once per-haplotype assembly with the same exceptions for the sex chromosomes as described above. The dipcall bed file was then curated to exclude reference regions and genomic alignments known to be problematic for benchmark generation. This includes portions of alignments with a break in the assembly-to-reference alignment inside a segmental duplication, a tandem repeat longer than 10 kbp, or a satellite repeat. We also excluded all gaps and homopolymers longer than 30 bp, since assembly-based variant calls are less accurate in these regions. Both high confidence bed files were then intersected to produce a final set of genomic regions which represent those confidently covered by both alignments of the super-scaffolded HPRC HG002 assembly. The TR catalog was then subset to TR regions contained entirely within the boundaries of a confidently covered region. All bed file operations were performed using BEDtools v2.30.0^70^.

The super-scaffolded HPRC HG002 assembly was then compared with Truvari v4.1 benchmarking and refinement to each alignment replicate as well as the Garg and Ebert HG002 technical replicates. To compare the benchmark to the technical replicates, Truvari refine was run with each sample’s bed file of confidently covered regions. The Truvari refine per-region bed file, which contains the initial benchmark regions’ coordinates and evaluation state (i.e. true positive, true negative, false negative, or false positive) were then joined across the comparison to the alignment and each technical replicate. Regions with a true positive or true negative state from comparison to the alignment replicate as well as at least one of the technical replicates were classified as being Tier1 and those failing this condition were classified as Tier2. Furthermore, sites where the state flipped between replicates (e.g. TP in the alignment replicate but TN in a technical replicate) were assumed to be unreliably compared and therefore demoted to Tier2. The union of the three replicates’ state was also recorded as an annotation column in the benchmark bed (e.g. TP_FN_FP corresponds to a true positive in the alignment replicate, false negative in the Ebert technical replicate, and false positive in the Garg technical replicate.

More annotations were then added to the benchmarking regions. First, the variants from the pVCF were analyzed for each benchmarking region in order to populate a ‘variant flag’. This flag uses bit-encoding to indicate if HG002 has a variant ≥5bp (bit 0×1), HG002 has a variant <5bp (0×2), another non-HG002 sample in the pVCF has a variant ≥5bp (bit 0×4), or a non-HG002 sample in the pVCF has a variant <5bp (bit 0×8). Additionally when analyzing the pVCF we summed the length of non-SNV variants for the maternal and paternal haplotypes in HG002 to record the region’s allele deltas. For example, a heterozygous 5bp deletion on the maternal allele and a second homozygous 5bp deletion would create a maternal allele delta of 10bp and a paternal allele delta of 5bp. Finally, the Shannon entropy of sequence spanned by the benchmarking region was calculated^79^ for identification of low-complexity sequences.

### Description of benchmarking software

Benchmarking results created using RTG vcfeval (https://github.com/RealTimeGenomics/rtg-tools) used version 3.12.1. Benchmarking results created using Truvari^30^ used version 4.1. While Truvari’s core variant comparison approach was previously published, for this project three additional features were developed to address the challenges of comparing variants between 5bp and 50bp, harmonizing variant representations, and generating performance reports for harmonized variants as well as the TR regions.

To handle comparison of variants as small as 5bp, the previously described^30^ reference context sequence similarity approach was replaced with an ‘unroll’ sequence similarity technique. The unroll technique works by first extracting the deleted reference sequence or inserted alternate sequence from two variants which we’ll call varA and varB. Next, the difference in start positions (posDiff) is calculated by subtracting varB’s start position from varA’s start position. If the posDiff is negative varA and varB are swapped and the absolute value of posDiff is taken. The number of base pairs that need to be ‘unrolled’ from varB (urLen) is calculated as the posDiff modulo the length of varB. The first urLen base pairs of varB’s sequence is then swapped with its remaining sequence. This is effectively equivalent to iteratively moving varB upstream by posDiff base pairs and swapping varB’s 5’ most base to the 3’ position of the sequence for each base moved. For example, imagine we have a reference sequence of ‘ATC’. We could have an expansion of this motif represented by an insertion at the 0th position with an alternate sequence of ‘ATC’, or represented as an insertion at the 2nd position of ‘CAT’; both of these variants would create a haplotype of ATCATC. The posDiff of these two variants is 2 and the urLen is 2 % 3 = 1, therefore moving the second variant’s first base C to create ‘ATC’. The edit distance is measured using edlib^80^ and is then converted into the longest common subsequence normalized sequence similarity using formula 1 - edit_distance(varA, varB) / (len(varA) + len(varB)).

Variant harmonization can be performed with a new Truvari command named ‘phab’. Inputs to phab include a VCF with phased variants, the reference against which the variants were called, and a set of regions to be harmonized. Using pysam, a python binding of htslib^81^, consensus sequences for haplotype 1 and haplotype 2 are created for each region. Each region’s haplotypes and reference sequence are then sent to MAFFT^82^ to perform multiple sequence alignment (MSA). MAFFT parameters are automatically chosen using their available --auto option. Alternatively, haplotypes may be independently aligned to their reference sequence with pyWFA, a cpython binding of the wave front aligner for harmonization^61^ (https://github.com/kcleal/pywfa). Truvari then converts the resultant MAFFT MSA or pyWFA pairwise alignments CIGAR strings back to VCF format.

Because some TR regions may not benefit from variant harmonization (e.g. some regions may have no variants from the benchmark, comparison callset, or either) It is unnecessary to run phab on all regions. We automate the selection of TR regions which could benefit from variant harmonization using another new Truvari command named ‘refine’. As input, refine takes a reference and a result from Truvari bench, which is a directory containing VCFs for true positive baseline, true positive comparison, false negative, and false positive variants as well as summary statistics and parameters provided to the call to Truvari bench. For each benchmarking region, the VCFs are parsed and variants counted. Each region with at least one false positive and one false negative variant - and therefore could benefit from variant harmonization - is collected and sent to phab. Refine will then re-compare the phab output VCF holding the baseline and comparison samples’ harmonized variants using the Truvari bench parameters initially provided by the user. For some benchmarking experiments, it may be beneficial to recreate haplotypes over all variants in the baseline/comparison VCFs (specifically single nucleotide variants and indels under the 5bp) instead of just the filtered set of variants contained in the initial Truvari bench output. Truvari refine provides a parameter --use-original-vcfs which will automatically pull variants from the original VCF files instead of the consolidation of benchmark result VCFs. One may want to use the original VCFs when recreating haplotypes where small variants would make more accurate haplotypes and would improve harmonization. Finally, the phab harmonization’s bench result is consolidated with the variant counts from regions which weren’t refined to create the final performance metric summary.

Since the Truvari phab variant harmonization process may alter variant counts, we also added a new reporting to the Truvari refine output which writes each benchmarking bed region with additional columns containing true positive, false negative, etc variant counts before and after Truvari phab was run. Each region is then assigned a true positive state if all variants in the region are matched between the baseline and comparison VCFs. If there are no variants in the region from either the baseline or comparison VCFs, the region is assigned a state of true negative. If there are any unmatched variants from the baseline, comparison, or both VCFs, the region is assigned a state of false negative, false positive, or false negative and false positive, respectively. These region state counts are then summarized and per-region metrics calculated. The set of per-region metrics and their definitions are provided in Supplementary Table 10.

To facilitate stratification analysis on Truvari results, we created a separate tool named Laytr (https://github.com/ACEnglish/laytr). Laytr will join the above described per-region report with the original TR benchmark bed file and the full TR catalog. This master table of region states and all available annotations described above is then grouped by various columns in the table and per-region metrics recalculated. Additionally, Laytr will take the self-organizing map which was built for this publication and is distributed with the benchmark as input and will build the hexagonal heatmaps similar to those found in Figure1 with the balanced accuracy per-neuron, accuracy per-neuron, and f1 per-neuron. The stratifications analyzed and reported are classified into eight sections: Benchmark subsets; Entropy; Gene TRs; Interspersed Repeats; Repeat complexity; Motif length; SOMs; Reference Expansion Contraction Mixed; Max Sizebin. All of these stratifications are recorded into a static html file which can be viewed in any web browser. Each stratification comes with a tooltip which documents in detail the definition of each section. In total, the eight sections provide 92 stratifications and the report provides 13 plots.

### Simulation of Tandem Repeats

To measure the difference of Truvari’s original reference context sequence similarity method with the new unroll sequence similarity method, we simulated 12,101,812 unique TR expansions with randomly generated motifs between 2bp and 20bp in length and between 2 and 50 copies of the motif. Each simulated TR is then expanded by between 1 and 15 copies and appended to the beginning of the TR to serve as the first alternate representation. This TR expansion then has a random selection of 0% to 30% (in 5% increments) of its bases randomly substituted to a different nucleotide before being placed randomly into the TR at any position except the start. This substituted and moved allele becomes the second alternate representation. This resulted in TR expansions between 2bp and 266bp in length being placed between 1bp and 931bp apart. All pairs of representations were then deduplicated before pairs were compared with the reference context, direct, and unroll sequence similarity methods before being contrasted with the simulated sequence similarity.

To test the functionality of Truvari phab variant harmonization approach, we slightly altered the above described TR simulation technique. Instead of a random motif, we chose motifs from the TR catalog from regions with above 95% mean motif purity. And, instead of randomly placing the second alternate representation in a single position downstream, we split it into between two and five subsequences before inserting each at different, consecutive positions. The two representations along with the reference sequence were then run through MAFFT^82^. Additionally, the two representations were each aligned to the reference sequence using a cpython binding of wave front aligner^61^ for harmonization (https://github.com/kcleal/pywfa). For both harmonization results per-simulation, the number of variant positions before and after harmonization were recorded and subsequently summarized.

### Generation of Tandem Repeat Callsets

To evaluate the utility of the TR benchmark we compared it to five tandem repeat callsets using Truvari. The four callsets were generated by TR calling tools: TRGT^20^; Medakka (https://github.com/nanoporetech/medaka); HipSTR^18^; GangSTR^19^. Additionally the GIAB v4.2.1 small variant benchmark^27^ was collected. Details of each callset’s generation are below. Callsets were compared using Truvari v4.1 with bench parameters comparing variants down to 5bp in length and each VCF entry was allowed to participate in as many matches as its allele count (e.g. a homozygous variant is allowed to match to a single homozygous variant or two heterozygous variants). Next, callsets were processed with Truvari refine using the original VCFs. The intersection of the refined regions’ states was generated and analyzed using custom scripts.

#### TRGT

TRGT v0.4.0 was run on a PacBio HiFi HG002 sample (available here https://www.pacb.com/connect/datasets/) with the default parameters. The adotto repeat catalog was reformatted to match TRGT input requirements. For repeat regions that corresponded to multiple motif sequences, we picked the motif sequence with the largest reference span.

#### Medaka

Medaka tandem (Medaka v1.9.0) was run with a custom model optimized for human consensus (https://cdn.oxfordnanoportal.com/software/analysis/models/medaka/res_medaka_tandem_r1041_e82_400bps_sup_v420.tar.gz) and with the adotto repeat catalog bed file given as “--regions”. Other parameters were left at defaults. The program was run on a 60× kit14 5kHz HG002 dataset basecalled with Dorado v0.2.5 with the v4.2.0 SUP basecalling model, haplotagged by Whatshap with SNPs called using Clair3.

#### HipSTR

We used HipSTR v0.7 with non-default parameters --def-stutter-model, --min-reads 15, and --haploid-chrs chrY,chrX to use default values for the stutter error model, consider loci for genotyping with at least 15 overlapping reads, and to assume chrX and chrY as haploid chromosomes. HipSTR was run on Illumina HiSeq 2×250 reads for HG002 obtained from GIAB (https://ftp-trace.ncbi.nlm.nih.gov/ReferenceSamples/giab/data/AshkenazimTrio/HG002_NA24385_son/NIST_Illumina_2X250bps/novoalign_bams/HG002.GRCh38.2X250.bam) to perform STR genotyping. The reference set of repeats were generated by subsetting this project’s TR catalog v1.1 to those in the HG002 TR benchmark. Regions with multiple repeat annotations were split into separate bed regions. Finally, all repeats with motif length above 6bp were excluded, resulting in a reference set of 1,826,383 repeats. We then used bcftools v1.12 norm-m-any to split multi-allelic records in HipSTR output VCF file, and ran Truvari to compare HipSTR records with benchmark. Truvari refine was run on the subset of 1,422,786 loci that had been successfully genotyped by HipSTR using the --regions parameter.

#### GangSTR

GangSTR results were generated using version 2.5.0 and processing the 60× HG002 bam aligned to GRCh38 and provided by NIST (https://ftp.ncbi.nlm.nih.gov/ReferenceSamples/giab/data/AshkenazimTrio/HG002_NA24385_son/NIST_HiSeq_HG002_Homogeneity-10953946/NHGRI_Illumina300X_AJtrio_novoalign_bams/HG002.GRCh38.60x.1.bam). The catalog of reference TR loci to analyze was generated from the full TR catalog^71^ by extracting the TandemRepeatsFinder^38^ position, period, and motif annotations (N=2,934,805). The resulting variants were then filtered to exclude calls with read depth below 15 and posterior probability Q score below 0.85. This filtering removed 221,074 (49.3%) variants. VCF was then processed by bcftools v1.12 norm to split multi-allelics and normalize indels to produce 227,167 variants which were then processed by Truvari as described above.

#### GIAB v4.2.1 small variant benchmark

The GIAB HG002 GRCh38 v4.2.1 small variant benchmark was downloaded from https://ftp-trace.ncbi.nlm.nih.gov/ReferenceSamples/giab/release/AshkenazimTrio/HG002_NA24385_son/NISTv4.2.1/GRCh38/HG002_GRCh38_1_22_v4.2.1_benchmark.vcf.gz. Prior to use in evaluations multi-allelic variants were normalized using ‘bcftools norm -m-any’ (bcftools v1.15). The output VCF was compressed and indexed with ‘bgzip’ and ‘tabix’ (htslib v1.14).

#### WGS Callers

The 60x HG002 bam aligned to GRCh38 and provided by NIST (https://ftp.ncbi.nlm.nih.gov/ReferenceSamples/giab/data/AshkenazimTrio/HG002_NA24385_son/NIST_HiSeq_HG002_Homogeneity-10953946/NHGRI_Illumina300X_AJtrio_novoalign_bams/HG002.GRCh38.60x.1.bam) was downloaded and used by two short-read WGS variant callers, DeepVariant version 1.4.0^83^ to produce SNPs and small indels, and BioGraph^84^ version 7.1.1. DeepVariant results were then processed by bcftools v1.12 norm to split multi-allelics and normalize indels. The long-read WGS caller Sniffles^62^ version 2.05 with min 50bp SV length and –phase option, was run on as part of PRINCESS pipeline^85^ (version 2) based on the phased bam file from GIAB PacBio HiFi 15kbp data set: https://ftp-trace.ncbi.nlm.nih.gov/ReferenceSamples/giab/data/AshkenazimTrio/HG002_NA24385_son/PacBio_CCS_15kb/. Each resulting VCF’s variants were then processed by Truvari as described above to compare to the benchmark.

## Data availability

The TR catalog v1.2 can be found at https://zenodo.org/records/8387564. Supplementary Table 4 holds the paths to the input assemblies used to create the pVCF. The pVCF can be found at https://zenodo.org/records/6975244. The TandemRepeat benchmark is hosted at https://ftp-trace.ncbi.nlm.nih.gov/ReferenceSamples/giab/release/AshkenazimTrio/HG002_NA24385_son/TandemRepeats_v1.0/

## Code availability

All code created for this project is available under an open-source license. Analysis scripts for this project are hosted at https://github.com/ACEnglish/adotto/. Truvari can be found at https://github.com/ACEnglish/truvari/. Laytr can be found at https://github.com/ACEnglish/laytr/. A lightweight version of the TR catalog creation process is available as a snakemake pipeline at https://github.com/nate-d-olson/adotto-smk. The overlap permutation tool regioners can be downloaded from https://github.com/ACEnglish/regioners.

## Acknowledgments

We would like to thank the GIAB community for constant support. We thank Jennifer McDaniel for very helpful comments on the manuscript. Mike Wykes and Sergey Nurk for assistance processing Medaka results. Vineet Bafna for contributions to the TR catalog. AE and FJS were supported by (HHSN268201800002I, U01 AG058589, 1U01HG011758-01, 1UG3NS132105-01). Helyaneh Ziaei-Jam supported by NIH/NHGRI R01HG010149. Mark Chaisson and Bida Gu supported by R01HG011649 and 5U24HG007497, respectively. Certain commercial equipment, instruments, or materials are identified to specify adequately experimental conditions or reported results. Such identification does not imply recommendation or endorsement by the National Institute of Standards and Technology, nor does it imply that the equipment, instruments, or materials identified are necessarily the best available for the purpose.

## Contributions

AE performed data analysis and software development. ED, HZ, NO, SM, JP, BG, JW, MG, MC contributed to testing and data processing. AE, JZ and FS designed the study. All the authors reviewed and edited the manuscript.

