## Supplementary Figures for "Benchmarking of small and large variants across tandem repeats"

### Supplementary Figure 1: Upset Plot

Upset Plot of each TR region source's contribution to the set of 2,171,789 candidate  TR regions. Combinations with fewer than 0.5% contribution not shown.

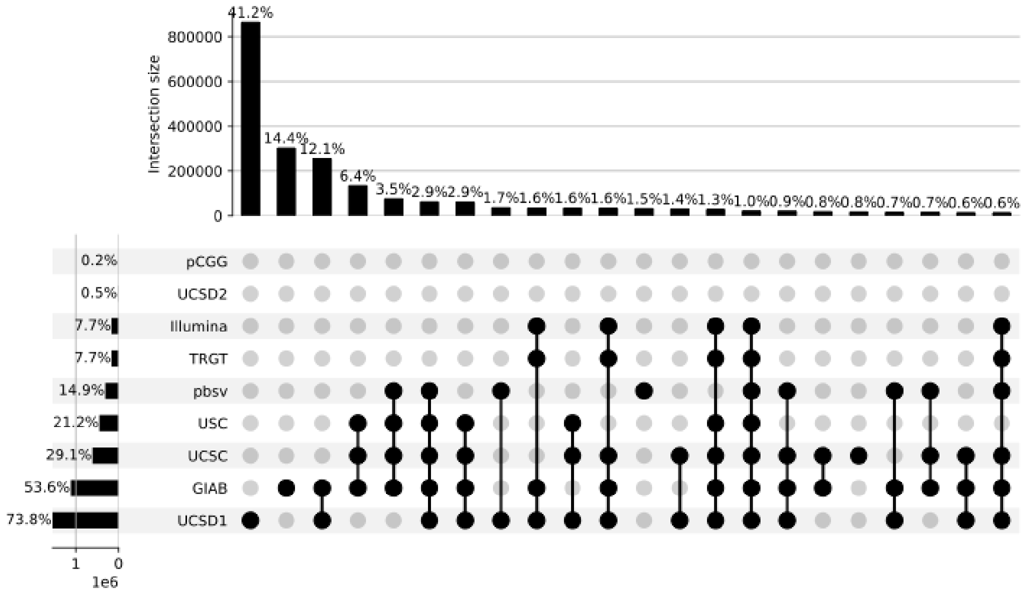

### Supplementary Figure 2: Buffer Length

TR catalog regions' upstream (blue/negative) and downstream (orange/positive) buffer length distribution. Input TR interval sources were expanded by ±25bp whereas the catalog's recorded buffer length is the distance between the region's start and end coordinates and the TandemRepeatsFinder first start and last end coordinates.

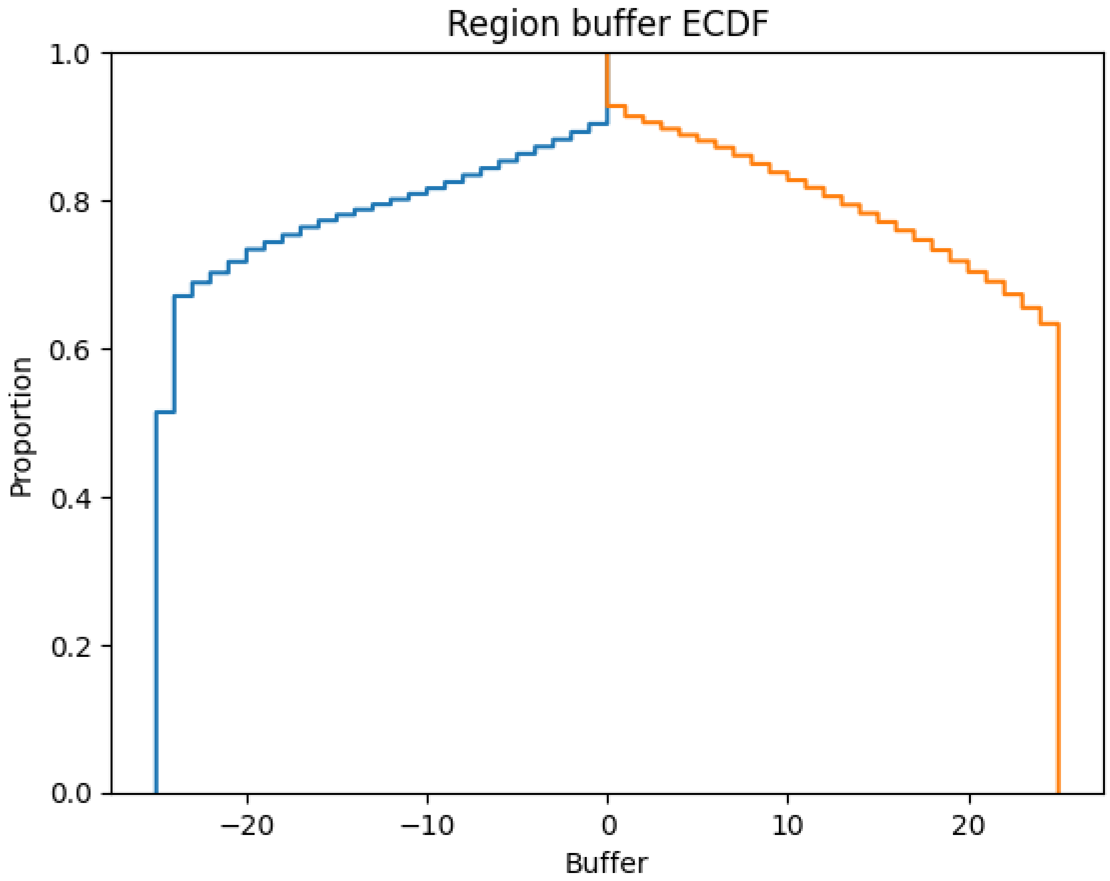

### Supplementary Figure 3: Catalog intersection

Intersection of TR catalog with promoters (top-left), transcripts (top-right), protein coding coding sequence (bottom-left) and protein coding untranslated regions (bottom-right). The distribution of the catalog's intersection to random permutations of intervals' coordinates are in gray. The observed intersections are marked with the vertical blue line.

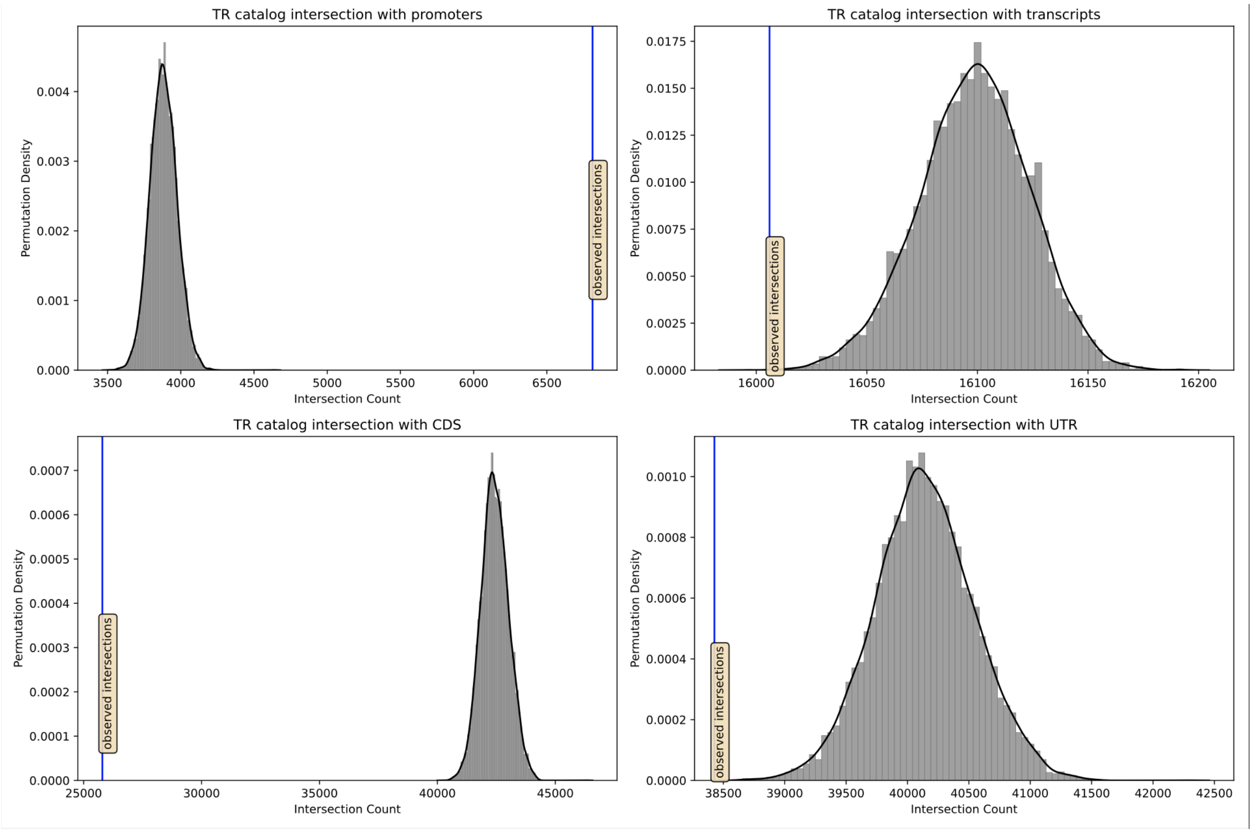

### Supplementary Figure 4: TR regions per-neuron

Distribution of the number of TR regions mapped to each SOM neuron.

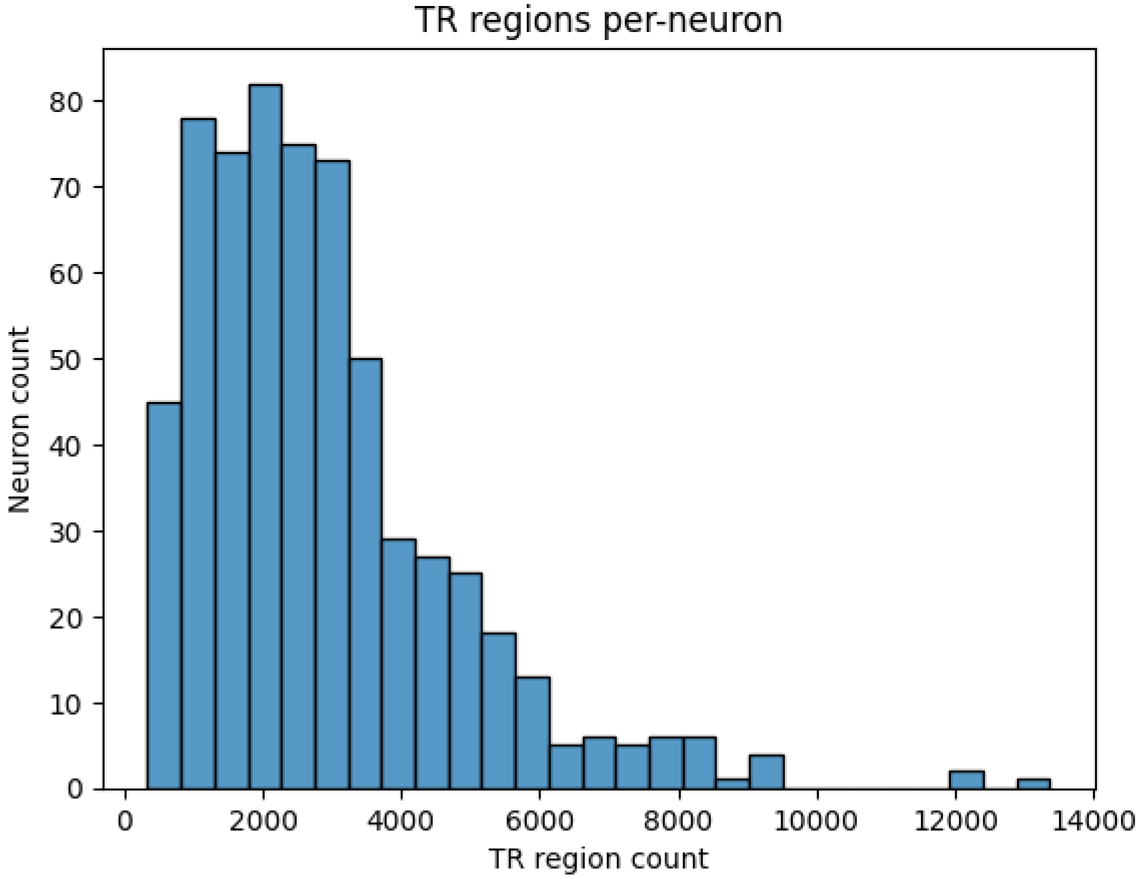

### Supplementary Figure 5: pVCF variant counts

Left - counts of pVCF variants. Right - proportion of pVCF variants with start and end positions within a single TR region. Each data points represents a single sample colored by sex.

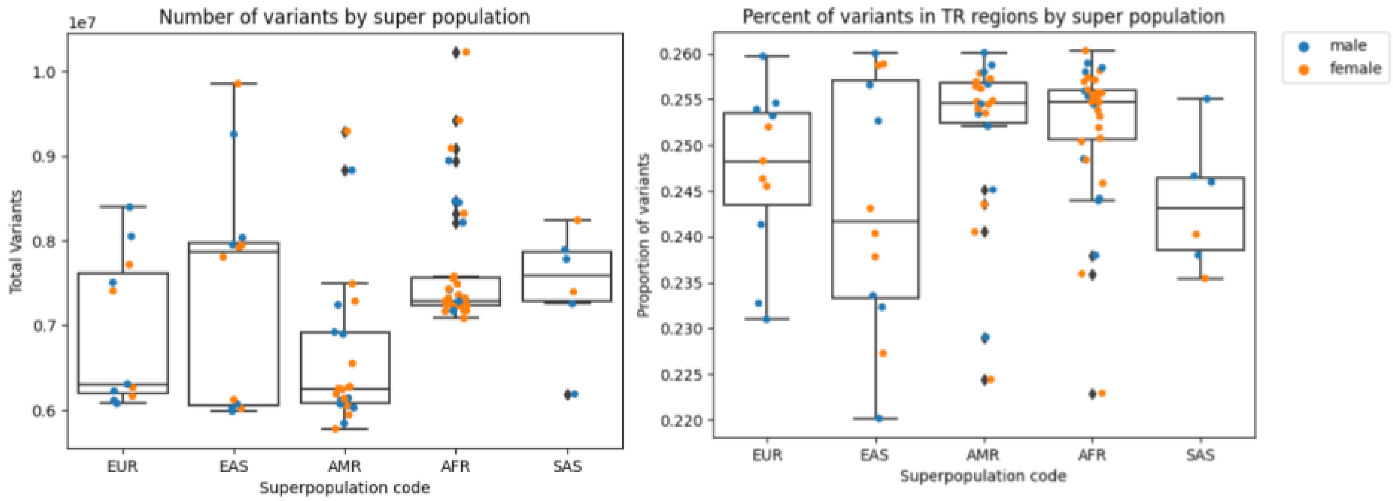

### Supplementary Figure 6: Neuron TR region percent in benchmark

Each TR region maps to a single neuron in the self-organizing map (SOM), The proportion of TR regions captured in the benchmark per neuron is calculated over the SOM's 625 neurons and distribution shown. Percent homopolymer for a TR region is calculated as the sum of bases annotated by TandemRepeatsFinder as homopolymer run divided by the total span of the TR region. Percent homopolymer for a SOM neuron is calculated as the mean of its TR regions' percent homopolymer.

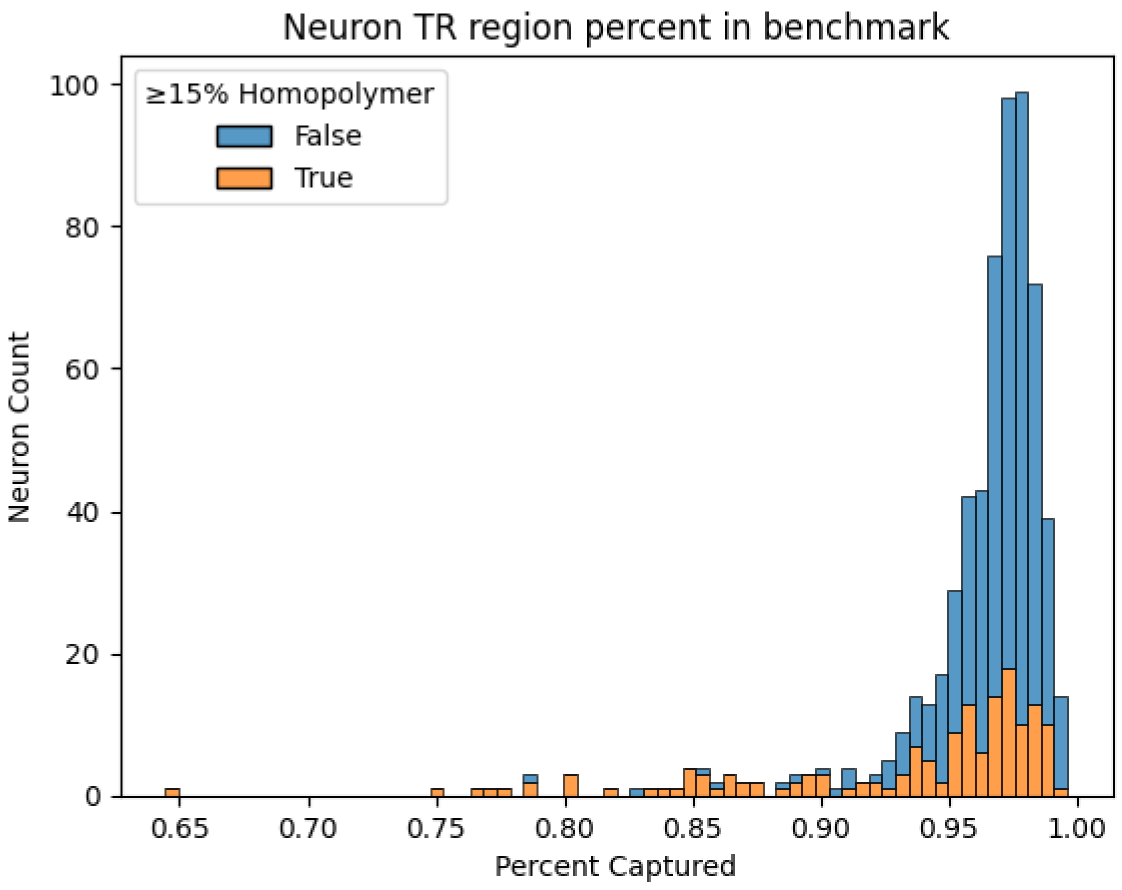

### Supplementary Figure 7: TR sequence similarity

For each simulated TR expansion (see methods), the sequences of the two representations were compared using Truvari's reference context sequence similarity, Truvari's unroll sequence similarity, and direct sequence similarity comparison of the inserted nucleotides.

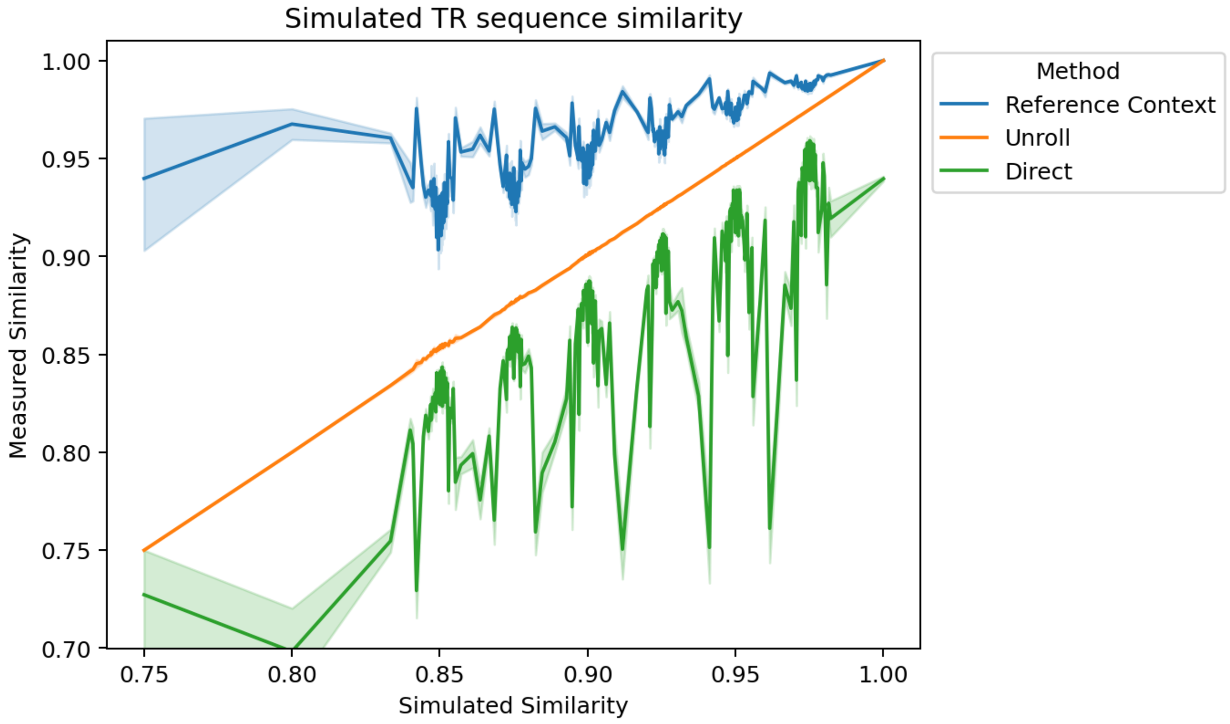
