## Supplementary Material 1 for "Benchmarking of small and large variants across tandem repeats"

### Benchmark subsets ?

Laytr

Download

| | TP | TN | FP | FN | base P | base N | comp P | comp N | PPV | TPR | TNR | NPV | ACC | BA | F1 |
|---|---|---|---|---|---|---|---|---|---|---|---|---|---|---|---|
| **Subset** | | | | | | | | | | | | | | | |
| **Full** | 103465 | 1594222 | 7418 | 2332 | 107253 | 1599600 | 111634 | 1595219 | 0.926823 | 0.964682 | 0.996638 | 0.999375 | 0.994630 | 0.980660 | 0.945374 |
| **Tier1** | 98934 | 1532505 | 5613 | 1743 | 101307 | 1537201 | 105147 | 1533361 | 0.940911 | 0.976576 | 0.996945 | 0.999442 | 0.995686 | 0.986761 | 0.958412 |
| **Tier2** | 4531 | 61717 | 1805 | 589 | 5946 | 62399 | 6487 | 61858 | 0.698474 | 0.762025 | 0.989070 | 0.997721 | 0.969317 | 0.875548 | 0.728867 |
| **AnyVar** | 103459 | 1115333 | 5997 | 2332 | 107247 | 1119289 | 110206 | 1116330 | 0.938778 | 0.964680 | 0.996466 | 0.999107 | 0.993686 | 0.980573 | 0.951553 |
| **>=5** | 103324 | 191841 | 4229 | 2321 | 107098 | 194027 | 108294 | 192831 | 0.954106 | 0.964761 | 0.988734 | 0.994866 | 0.980208 | 0.976747 | 0.959404 |

### Entropy ?

Download

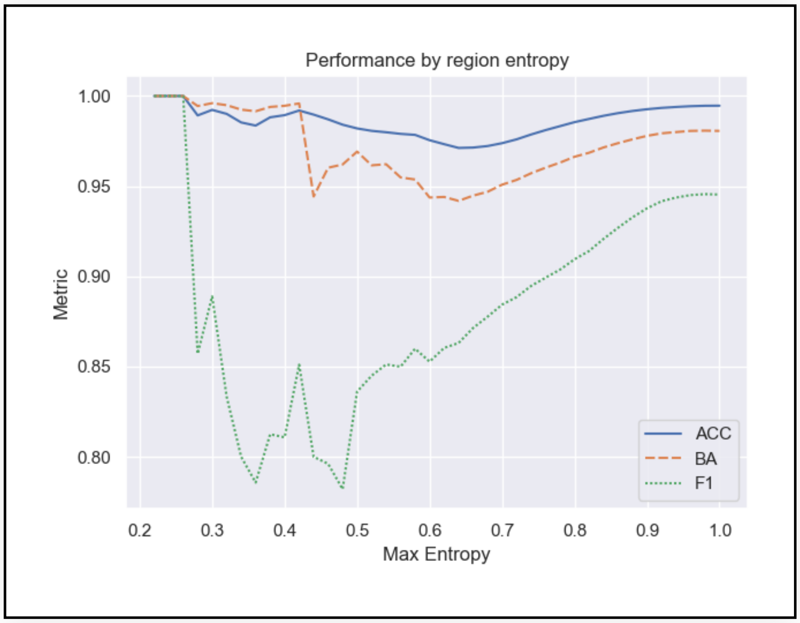


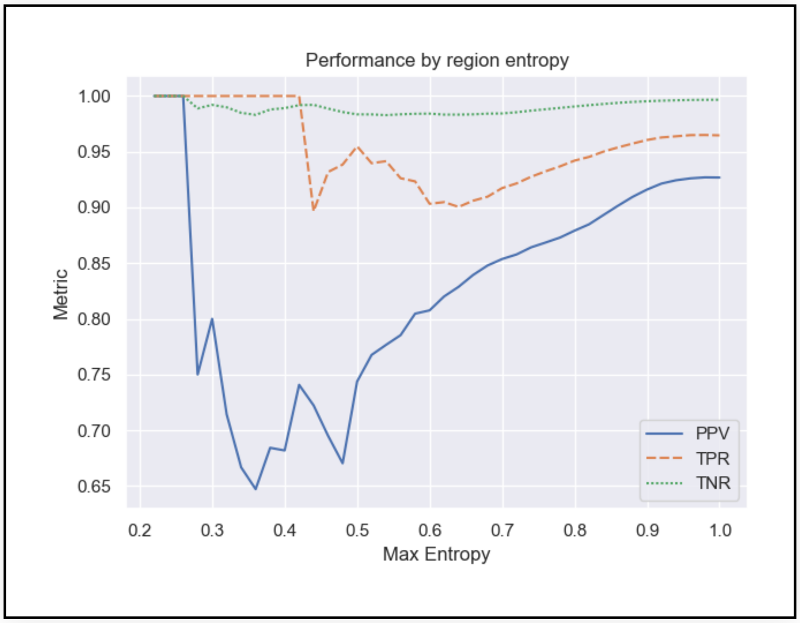


### Gene TRs ?

Download

| | TP | TN | FP | FN | base P | base N | comp P | comp N | PPV | TPR | TNR | NPV | ACC | BA | F1 |
|---|---|---|---|---|---|---|---|---|---|---|---|---|---|---|---|
| **Subset** | | | | | | | | | | | | | | | |
| **Intergenic** | 41244 | 595046 | 3023 | 970 | 42883 | 597144 | 44566 | 595461 | 0.925459 | 0.961780 | 0.996487 | 0.999303 | 0.994161 | 0.979133 | 0.943270 |
| **Genic** | 62221 | 999176 | 4395 | 1362 | 64370 | 1002456 | 67068 | 999758 | 0.927730 | 0.966615 | 0.996728 | 0.999418 | 0.994911 | 0.981671 | 0.946773 |
| **Protein Coding Gene** | 46002 | 756674 | 3265 | 975 | 47566 | 759121 | 49607 | 757080 | 0.927329 | 0.967119 | 0.996777 | 0.999464 | 0.995028 | 0.981948 | 0.946806 |

### Interspersed Repeats ?

Download

| | TP | TN | FP | FN | base P | base N | comp P | comp N | PPV | TPR | TNR | NPV | ACC | BA | F1 |
|---|---|---|---|---|---|---|---|---|---|---|---|---|---|---|---|
| **Subset** | | | | | | | | | | | | | | | |
| **No Interspersed** | 95420 | 1512099 | 6821 | 2077 | 98805 | 1517070 | 102919 | 1512956 | 0.927137 | 0.965741 | 0.996723 | 0.999434 | 0.994829 | 0.981232 | 0.946045 |
| **Interspersed** | 8045 | 82123 | 597 | 255 | 8448 | 82530 | 8715 | 82263 | 0.923121 | 0.952296 | 0.995068 | 0.998298 | 0.991097 | 0.973682 | 0.937482 |

### Repeat Complexity ?

Download

| | TP | TN | FP | FN | base P | base N | comp P | comp N | PPV | TPR | TNR | NPV | ACC | BA | F1 |
|---|---|---|---|---|---|---|---|---|---|---|---|---|---|---|---|
| **Subset** | | | | | | | | | | | | | | | |
| **Simple Overlap** | 41173 | 820774 | 3946 | 895 | 42583 | 824003 | 45435 | 821151 | 0.906196 | 0.966888 | 0.996081 | 0.999541 | 0.994647 | 0.981485 | 0.935559 |
| **Parent Overlap** | 14387 | 291046 | 940 | 310 | 14906 | 291673 | 15424 | 291155 | 0.932767 | 0.965182 | 0.997850 | 0.999626 | 0.996262 | 0.981516 | 0.948698 |
| **Complex Overlap** | 47905 | 482402 | 2532 | 1127 | 49764 | 483924 | 50775 | 482913 | 0.943476 | 0.962644 | 0.996855 | 0.998942 | 0.993665 | 0.979749 | 0.952964 |
| **Multi Anno** | 50277 | 517773 | 2650 | 1199 | 52240 | 519352 | 53282 | 518310 | 0.943602 | 0.962423 | 0.996960 | 0.998964 | 0.993803 | 0.979692 | 0.952920 |
| **Multi SubReg** | 44392 | 436341 | 2304 | 1055 | 46135 | 437702 | 47008 | 436829 | 0.944350 | 0.962220 | 0.996891 | 0.998883 | 0.993585 | 0.979555 | 0.953201 |

### Motif Length ?

Download

| | TP | TN | FP | FN | base P | base N | comp P | comp N | PPV | TPR | TNR | NPV | ACC | BA | F1 |
|---|---|---|---|---|---|---|---|---|---|---|---|---|---|---|---|
| **Motif Length Bin** | | | | | | | | | | | | | | | |
| **<5** | 55848 | 786785 | 4188 | 1233 | 57886 | 789882 | 60497 | 787271 | 0.923153 | 0.964793 | 0.996079 | 0.999383 | 0.993943 | 0.980436 | 0.943514 |
| **[5,10)** | 16688 | 344496 | 1239 | 358 | 17251 | 345476 | 18025 | 344702 | 0.925825 | 0.967364 | 0.997163 | 0.999402 | 0.995746 | 0.982264 | 0.946139 |
| **[10, 20)** | 12950 | 289699 | 961 | 274 | 13414 | 290393 | 13992 | 289815 | 0.925529 | 0.965409 | 0.997610 | 0.999600 | 0.996188 | 0.981510 | 0.945049 |
| **[20, 50)** | 10791 | 116982 | 563 | 229 | 11161 | 117319 | 11410 | 117070 | 0.945749 | 0.966849 | 0.997127 | 0.999248 | 0.994497 | 0.981988 | 0.956183 |
| **>=50** | 7188 | 56260 | 467 | 238 | 7541 | 56530 | 7710 | 56361 | 0.932296 | 0.953189 | 0.995224 | 0.998208 | 0.990276 | 0.974207 | 0.942627 |

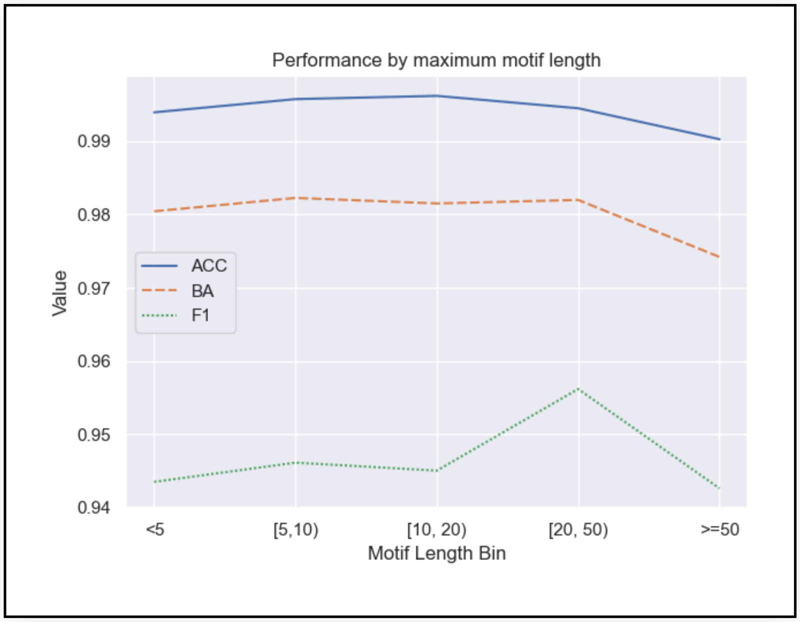


### Homopolymers ?

Download

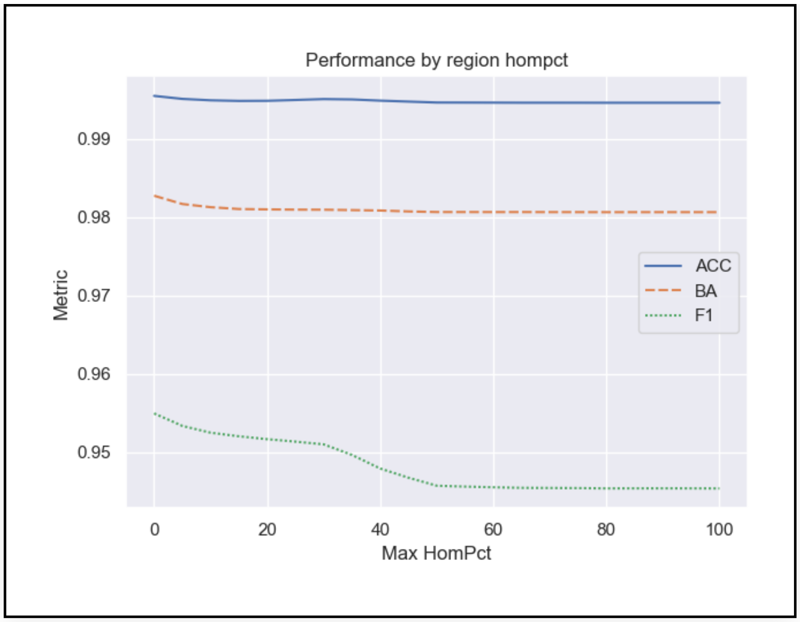
Performance by region hompct
Metric
Max HomPct
0.99
0.98
0.97
0.96
0.95
0
20
40
60
80
100
ACC
BA
F1

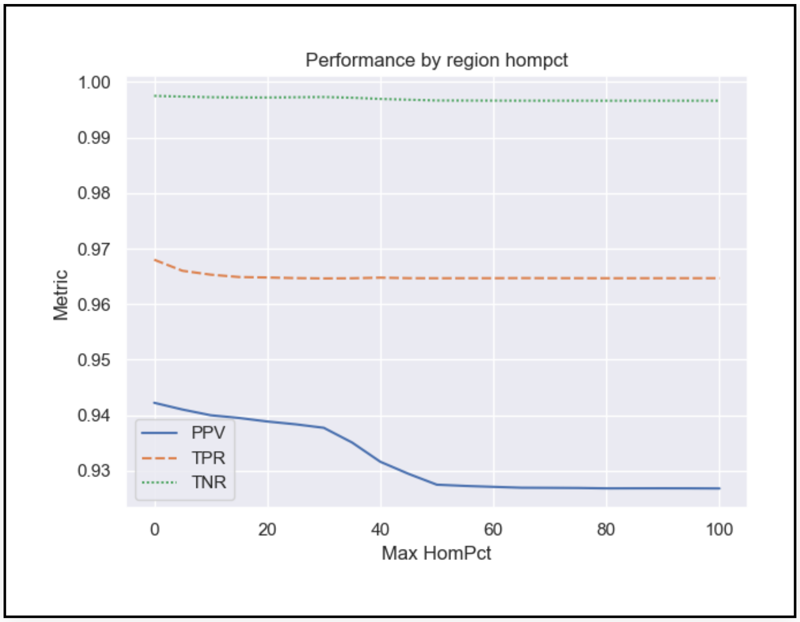
Performance by region hompct
Metric
Max HomPct
1.00
0.99
0.98
0.97
0.96
0.95
0.94
0.93
0
20
40
60
80
100
PPV
TPR
TNR

#### SOMs ?

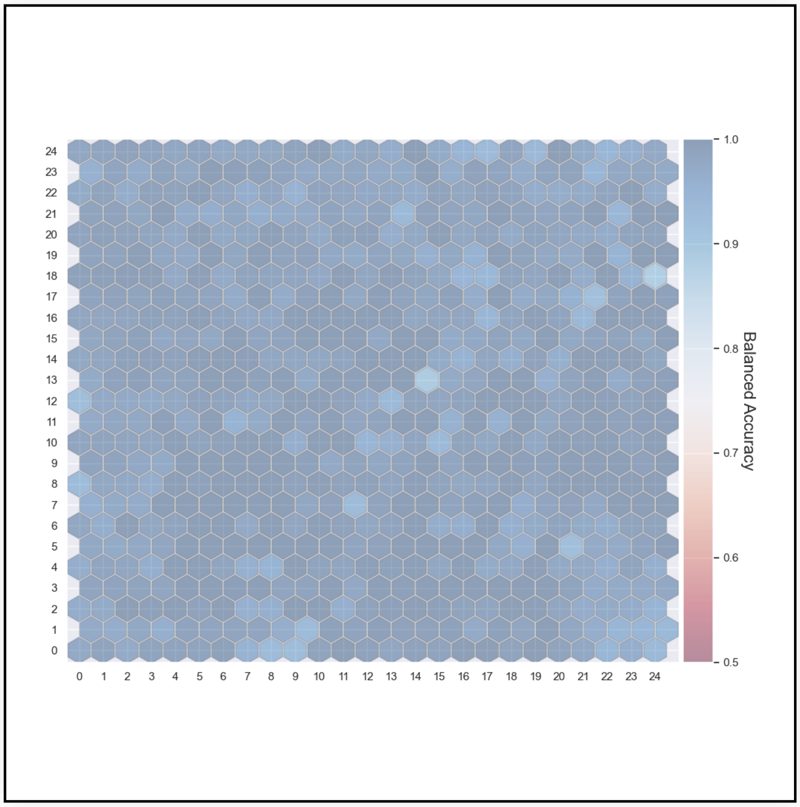
Balanced Accuracy
1.0
0.9
0.8
0.7
0.6
0.5

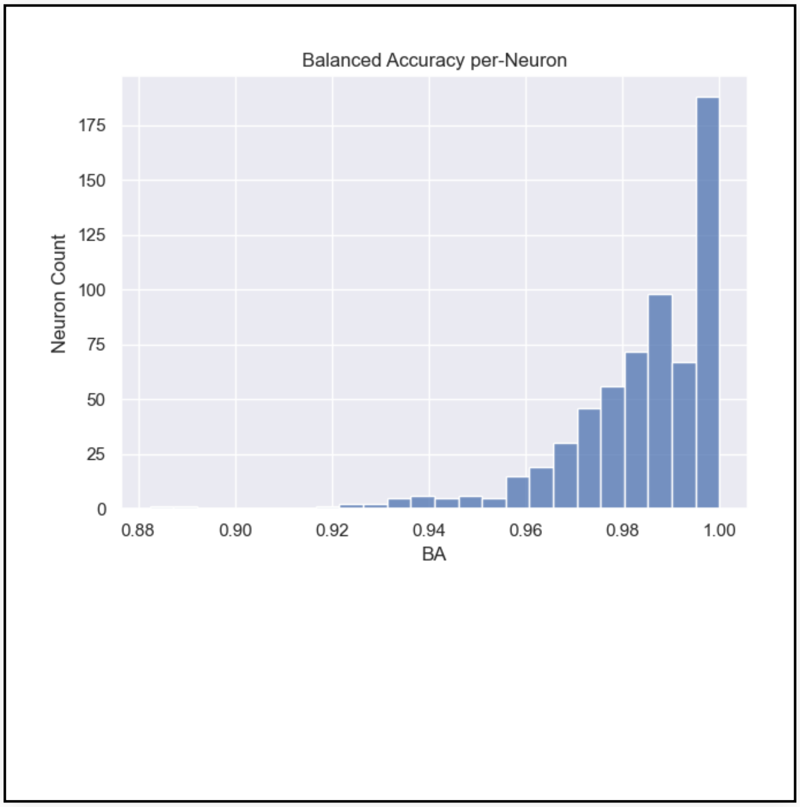
Balanced Accuracy per-Neuron
Neuron Count
BA
175
150
125
100
75
50
25
0
0.88
0.90
0.92
0.94
0.96
0.98
1.00

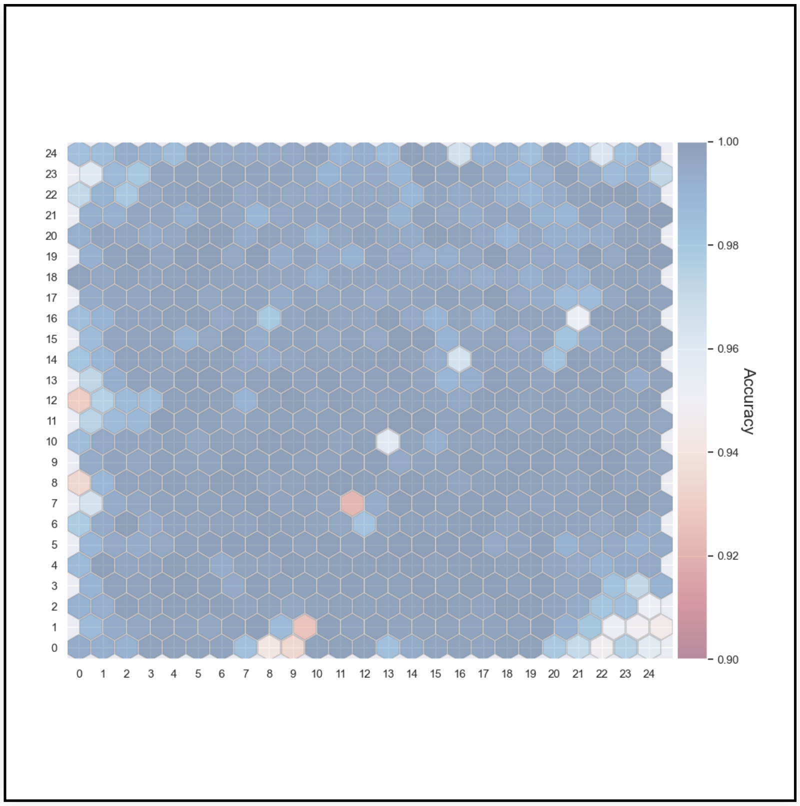


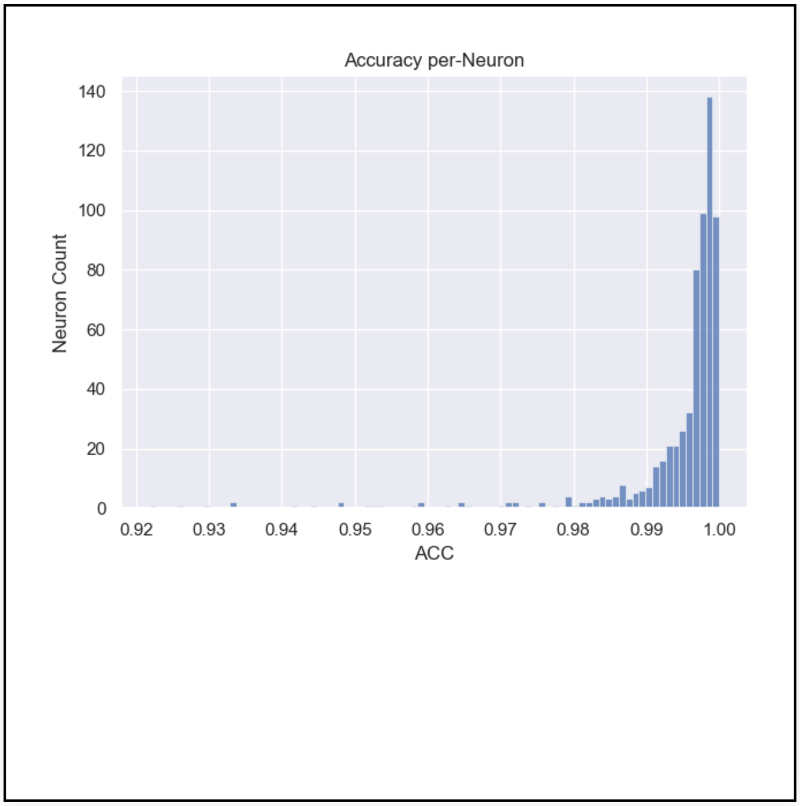


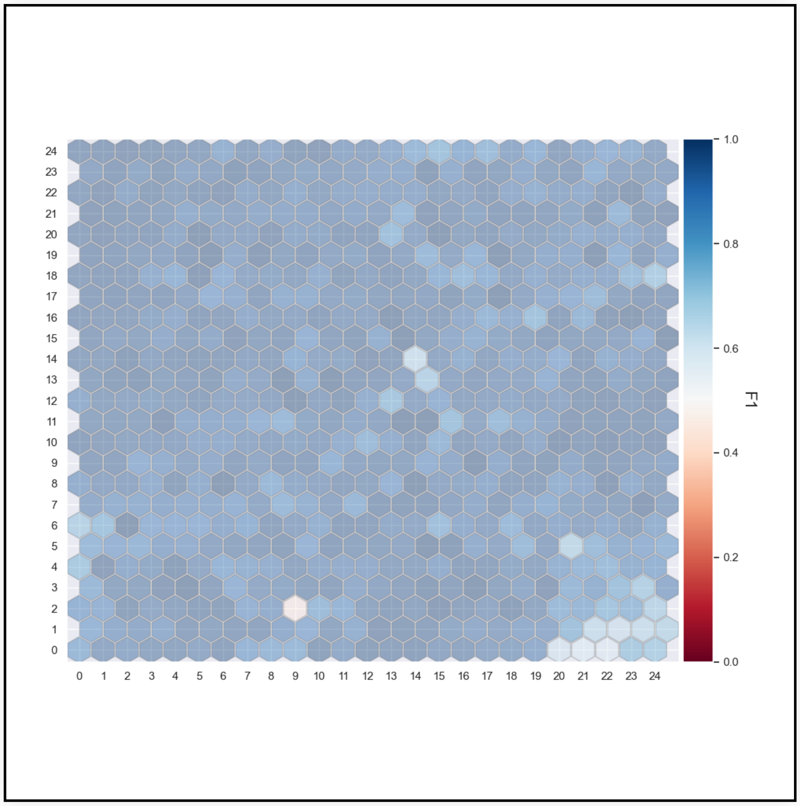


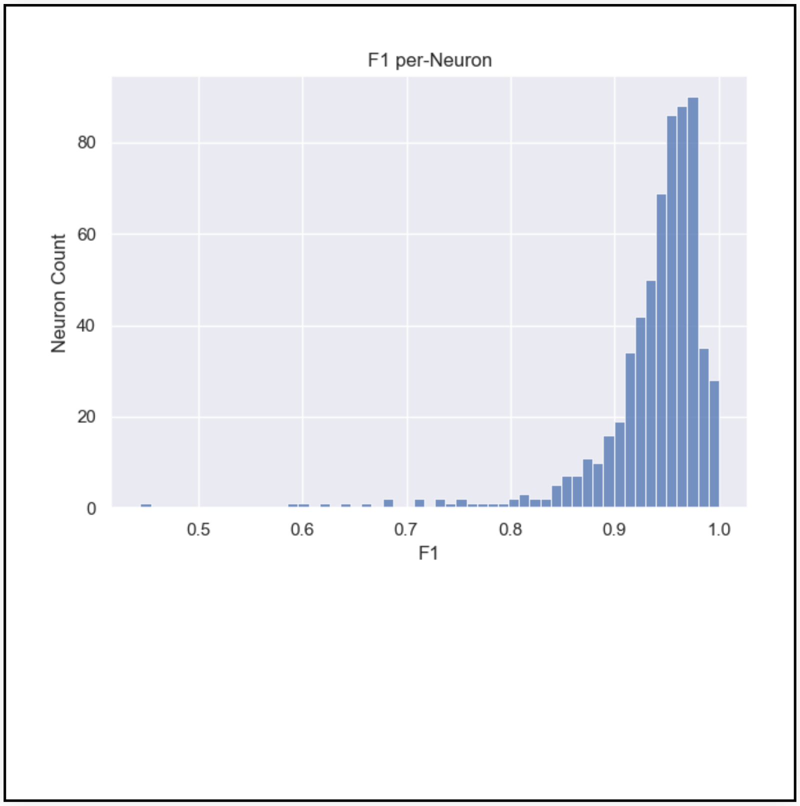


#### Reference Expansion Contraction Mixed* ?

Download

| | TP | TN | FP | FN | base P | base N | comp P | comp N | PPV | TPR | TNR | NPV | ACC | BA | F1 |
|---|---|---|---|---|---|---|---|---|---|---|---|---|---|---|---|
| ExpCon | | | | | | | | | | | | | | | |
| EXP | 103204 | 195806 | 3210 | 2316 | 106969 | 196982 | 107155 | 196796 | 0.963128 | 0.964803 | 0.994030 | 0.994969 | 0.983744 | 0.979416 | 0.963965 |
| REF | 261 | 1398416 | 4208 | 16 | 284 | 1402618 | 4479 | 1398423 | 0.058272 | 0.919014 | 0.997004 | 0.999995 | 0.996988 | 0.958009 | 0.109595 |

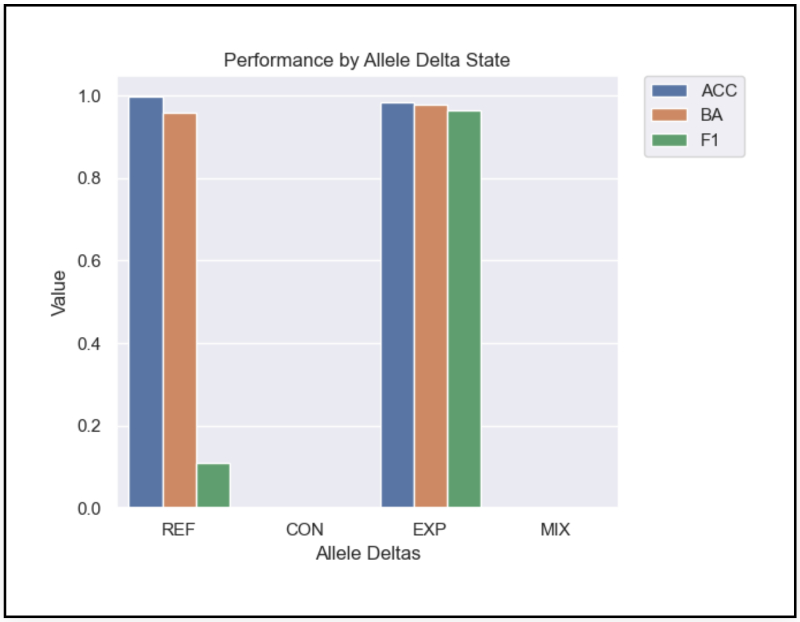


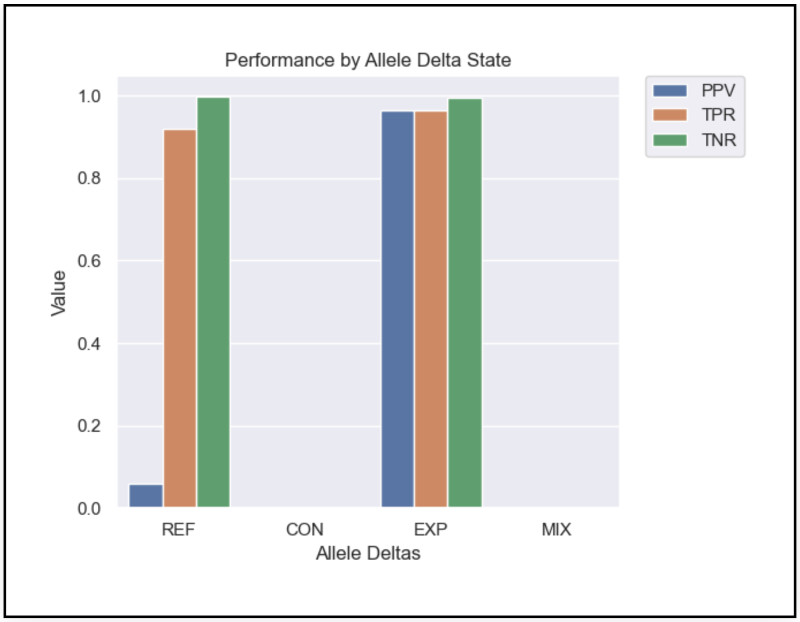


#### Max SizeBin* ?

Download

| | TP | TN | FP | FN | base P | base N | comp P | comp N | PPV | TPR | TNR | NPV | ACC | BA | F1 |
|---|---|---|---|---|---|---|---|---|---|---|---|---|---|---|---|
| **SNP** | 261 | 1398416 | 4208 | 16 | 284 | 1402618 | 4479 | 1398423 | 0.058272 | 0.919014 | 0.997004 | 0.999995 | 0.996988 | 0.958009 | 0.109595 |
| **[1,5)** | 221 | 191232 | 1061 | 20 | 252 | 192279 | 1293 | 191238 | 0.170920 | 0.876984 | 0.994555 | 0.999969 | 0.994401 | 0.935769 | 0.286084 |
| **[5,10)** | 39696 | 3990 | 635 | 959 | 41046 | 4095 | 40498 | 4643 | 0.980197 | 0.967110 | 0.974359 | 0.859358 | 0.967768 | 0.970735 | 0.973609 |
| **[10,15)** | 21872 | 278 | 406 | 398 | 22571 | 287 | 22462 | 396 | 0.973733 | 0.969031 | 0.968641 | 0.702020 | 0.969026 | 0.968836 | 0.971377 |
| **[15,20)** | 10377 | 59 | 189 | 178 | 10701 | 61 | 10662 | 100 | 0.973270 | 0.969722 | 0.967213 | 0.590000 | 0.969708 | 0.968468 | 0.971493 |
| **[20,30)** | 11379 | 58 | 242 | 207 | 11762 | 60 | 11724 | 98 | 0.970573 | 0.967438 | 0.966667 | 0.591837 | 0.967434 | 0.967052 | 0.969003 |
| **[30,40)** | 4997 | 21 | 120 | 99 | 5183 | 24 | 5159 | 48 | 0.968599 | 0.964113 | 0.875000 | 0.437500 | 0.963703 | 0.919557 | 0.966351 |
| **[40,50)** | 2588 | 13 | 63 | 47 | 2689 | 14 | 2675 | 28 | 0.967477 | 0.962440 | 0.928571 | 0.464286 | 0.962264 | 0.945505 | 0.964952 |
| **[50,100)** | 4834 | 10 | 94 | 73 | 4983 | 11 | 4964 | 30 | 0.973811 | 0.970098 | 0.909091 | 0.333333 | 0.969964 | 0.939595 | 0.971951 |
| **[100,200)** | 2959 | 17 | 64 | 44 | 3055 | 18 | 3041 | 32 | 0.973035 | 0.968576 | 0.944444 | 0.531250 | 0.968435 | 0.956510 | 0.970801 |
| **[200,300)** | 1161 | 5 | 32 | 21 | 1202 | 5 | 1197 | 10 | 0.969925 | 0.965890 | 1.000000 | 0.500000 | 0.966031 | 0.982945 | 0.967903 |
| **[300,400)** | 782 | 29 | 37 | 24 | 831 | 30 | 827 | 34 | 0.945586 | 0.941035 | 0.966667 | 0.852941 | 0.941928 | 0.953851 | 0.943305 |
| **[400,600)** | 744 | 42 | 48 | 31 | 805 | 43 | 800 | 48 | 0.930000 | 0.924224 | 0.976744 | 0.875000 | 0.926887 | 0.950484 | 0.927103 |
| **[600,800)** | 452 | 24 | 30 | 21 | 492 | 24 | 490 | 26 | 0.922449 | 0.918699 | 1.000000 | 0.923077 | 0.922481 | 0.959350 | 0.920570 |
| **[800,1k)** | 267 | 10 | 20 | 13 | 293 | 10 | 291 | 12 | 0.917526 | 0.911263 | 1.000000 | 0.833333 | 0.914191 | 0.955631 | 0.914384 |
| **[1k,2.5k)** | 670 | 14 | 97 | 80 | 782 | 16 | 778 | 20 | 0.861183 | 0.856777 | 0.875000 | 0.700000 | 0.857143 | 0.865889 | 0.858974 |
| **[2.5k,5k)** | 155 | 1 | 41 | 48 | 214 | 1 | 206 | 9 | 0.752427 | 0.724299 | 1.000000 | 0.111111 | 0.725581 | 0.862150 | 0.738095 |
| **>=5k** | 50 | 3 | 31 | 53 | 108 | 4 | 88 | 24 | 0.568182 | 0.462963 | 0.750000 | 0.125000 | 0.473214 | 0.606481 | 0.510204 |

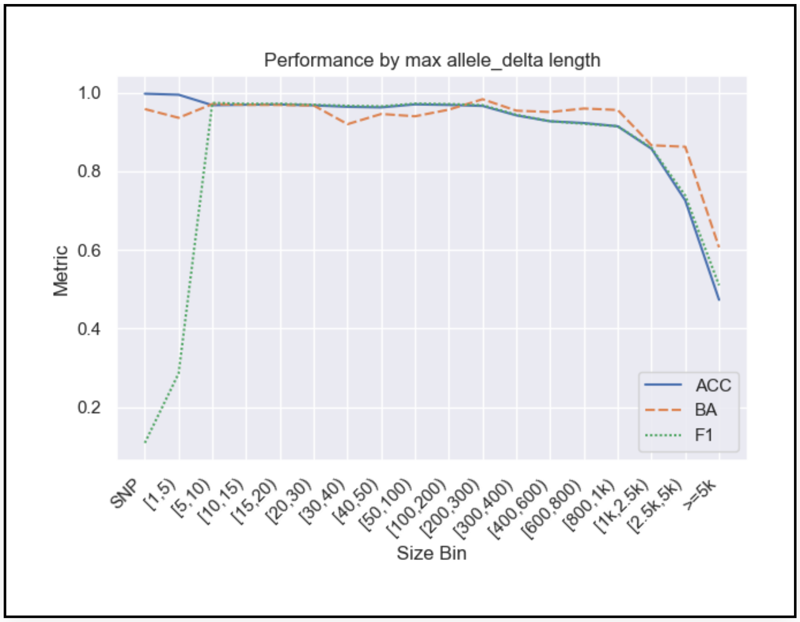


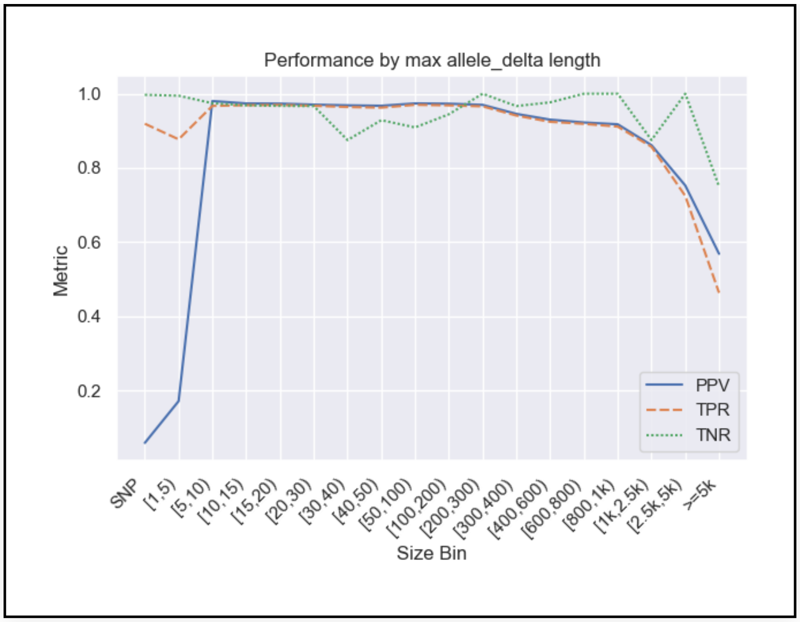


*Stratification by sizes requires a present variant. Therefore metrics which use TN (e.g. TNR, NPV, ACC) aren't informative.
