## Supplementary Material 2 for "Benchmarking of small and large variants across tandem repeats"

### Benchmark subsets ?

Download

| | TP | TN | FP | FN | base P | base N | comp P | comp N | PPV | TPR | TNR | NPV | ACC | BA | F1 |
|---|---|---|---|---|---|---|---|---|---|---|---|---|---|---|---|
| **Subset** | | | | | | | | | | | | | | | |
| **Full** | 7593 | 1601681 | 731 | 97424 | 105081 | 1601772 | 11949 | 1694904 | 0.635451 | 0.072259 | 0.999943 | 0.944998 | 0.942831 | 0.536101 | 0.129762 |
| **Tier1** | 7195 | 1536766 | 530 | 94425 | 101673 | 1536835 | 11070 | 1627438 | 0.649955 | 0.070766 | 0.999955 | 0.944285 | 0.942297 | 0.535361 | 0.127635 |
| **Tier2** | 398 | 64915 | 201 | 2999 | 3408 | 64937 | 879 | 67466 | 0.452787 | 0.116784 | 0.999661 | 0.962188 | 0.955637 | 0.558223 | 0.185678 |
| **AnyVar** | 7593 | 1121375 | 726 | 97418 | 105075 | 1121461 | 11944 | 1214592 | 0.635717 | 0.072263 | 0.999923 | 0.923252 | 0.920452 | 0.536093 | 0.129774 |
| **>=5** | 7589 | 196025 | 704 | 97381 | 105034 | 196091 | 11917 | 289208 | 0.636821 | 0.072253 | 0.999663 | 0.677799 | 0.676178 | 0.535958 | 0.129781 |

### Entropy ?

Download

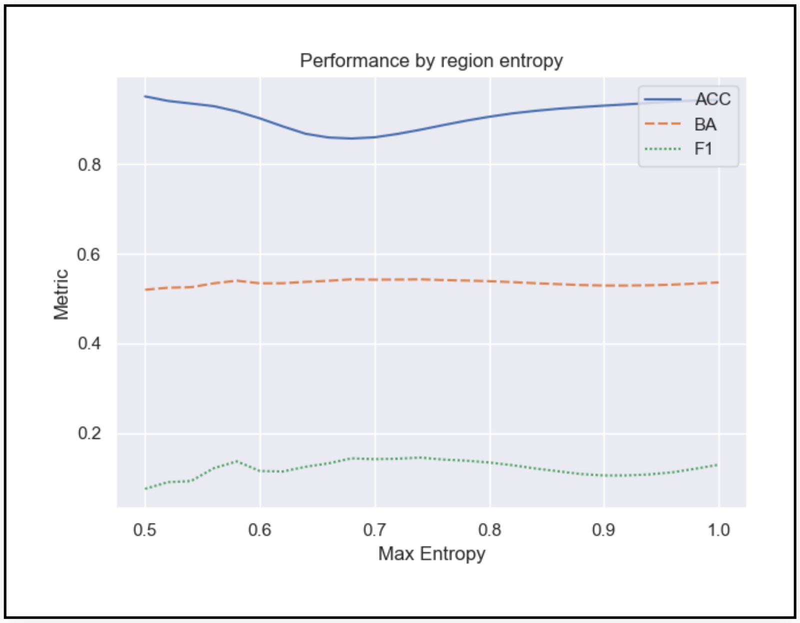


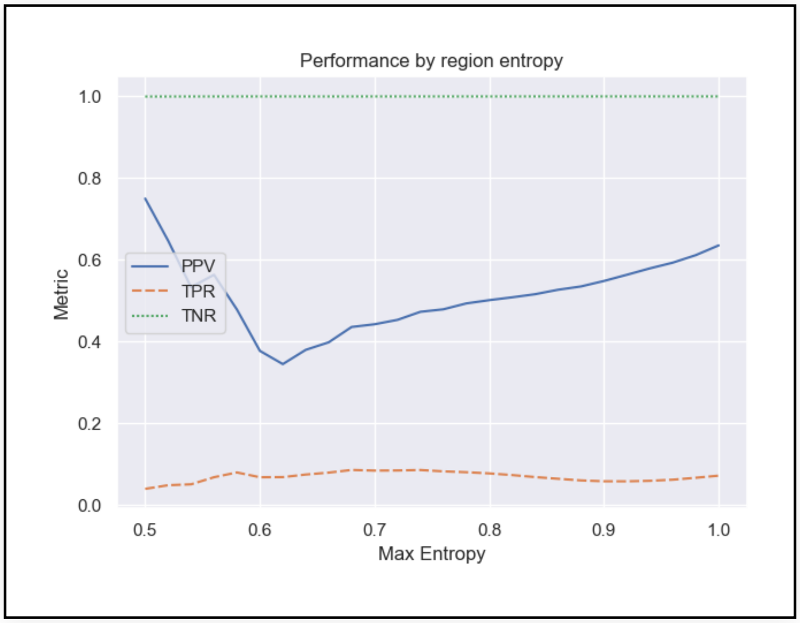


### Gene TRs ?

Download

| | TP | TN | FP | FN | base P | base N | comp P | comp N | PPV | TPR | TNR | NPV | ACC | BA | F1 |
|---|---|---|---|---|---|---|---|---|---|---|---|---|---|---|---|
| **Subset** | | | | | | | | | | | | | | | |
| **Intergenic** | 2918 | 598393 | 315 | 38643 | 41590 | 598437 | 4738 | 635289 | 0.615872 | 0.070161 | 0.999926 | 0.941922 | 0.939509 | 0.535044 | 0.125971 |
| **Genic** | 4675 | 1003288 | 416 | 58781 | 63491 | 1003335 | 7211 | 1059615 | 0.648315 | 0.073632 | 0.999953 | 0.946842 | 0.944824 | 0.536793 | 0.132245 |
| **Protein Coding Gene** | 3621 | 759711 | 320 | 43296 | 46944 | 759743 | 5522 | 801165 | 0.655741 | 0.077134 | 0.999958 | 0.948258 | 0.946255 | 0.538546 | 0.138032 |

### Interspersed Repeats ?

Download

| | TP | TN | FP | FN | base P | base N | comp P | comp N | PPV | TPR | TNR | NPV | ACC | BA | F1 |
|---|---|---|---|---|---|---|---|---|---|---|---|---|---|---|---|
| **Subset** | | | | | | | | | | | | | | | |
| **No Interspersed** | 6740 | 1519089 | 684 | 89913 | 96710 | 1519165 | 10788 | 1605087 | 0.624768 | 0.069693 | 0.999950 | 0.946422 | 0.944274 | 0.534821 | 0.125398 |
| **Interspersed** | 853 | 82592 | 47 | 7511 | 8371 | 82607 | 1161 | 89817 | 0.734711 | 0.101899 | 0.999818 | 0.919559 | 0.917200 | 0.550859 | 0.178976 |

### Repeat Complexity ?

Download

| | TP | TN | FP | FN | base P | base N | comp P | comp N | PPV | TPR | TNR | NPV | ACC | BA | F1 |
|---|---|---|---|---|---|---|---|---|---|---|---|---|---|---|---|
| **Subset** | | | | | | | | | | | | | | | |
| **Simple Overlap** | 1549 | 824956 | 112 | 40051 | 41607 | 824979 | 2300 | 864286 | 0.673478 | 0.037229 | 0.999972 | 0.954494 | 0.953748 | 0.518601 | 0.070558 |
| **Parent Overlap** | 2267 | 291915 | 242 | 12355 | 14642 | 291937 | 3501 | 303078 | 0.647529 | 0.154829 | 0.999925 | 0.963168 | 0.959563 | 0.577377 | 0.249904 |
| **Complex Overlap** | 3777 | 484810 | 377 | 45018 | 48832 | 484856 | 6148 | 527540 | 0.614346 | 0.077347 | 0.999905 | 0.919001 | 0.915492 | 0.538626 | 0.137395 |
| **Multi Anno** | 4668 | 520240 | 480 | 46590 | 51300 | 520292 | 7365 | 564227 | 0.633809 | 0.090994 | 0.999900 | 0.922040 | 0.918326 | 0.545447 | 0.159141 |
| **Multi SubReg** | 3096 | 438543 | 310 | 42128 | 45255 | 438582 | 5141 | 478696 | 0.602217 | 0.068412 | 0.999911 | 0.916120 | 0.912785 | 0.534162 | 0.122867 |

### Motif Length ?

Download

| | TP | TN | FP | FN | base P | base N | comp P | comp N | PPV | TPR | TNR | NPV | ACC | BA | F1 |
|---|---|---|---|---|---|---|---|---|---|---|---|---|---|---|---|
| **Motif Length Bin** | | | | | | | | | | | | | | | |
| **<5** | 1234 | 791273 | 77 | 55239 | 56481 | 791287 | 2173 | 845595 | 0.567879 | 0.021848 | 0.999982 | 0.935759 | 0.934816 | 0.510915 | 0.042077 |
| **[5,10)** | 635 | 345809 | 49 | 16270 | 16910 | 345817 | 1116 | 361611 | 0.568996 | 0.037552 | 0.999977 | 0.956301 | 0.955109 | 0.518764 | 0.070454 |
| **[10, 20)** | 855 | 290598 | 85 | 12336 | 13195 | 290612 | 1528 | 302279 | 0.559555 | 0.064797 | 0.999952 | 0.961357 | 0.959336 | 0.532375 | 0.116145 |
| **[20, 50)** | 2222 | 117453 | 321 | 8754 | 11003 | 117477 | 3793 | 124687 | 0.585816 | 0.201945 | 0.999796 | 0.941983 | 0.931468 | 0.600870 | 0.300351 |
| **>=50** | 2647 | 56548 | 199 | 4825 | 7492 | 56579 | 3339 | 60732 | 0.792752 | 0.353310 | 0.999452 | 0.931107 | 0.923897 | 0.676381 | 0.488782 |

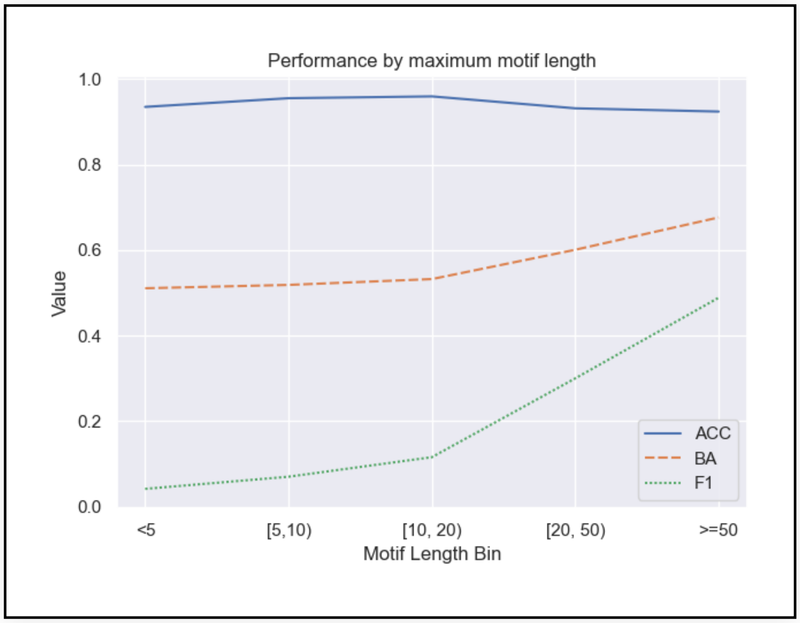


### Homopolymers ?

Download

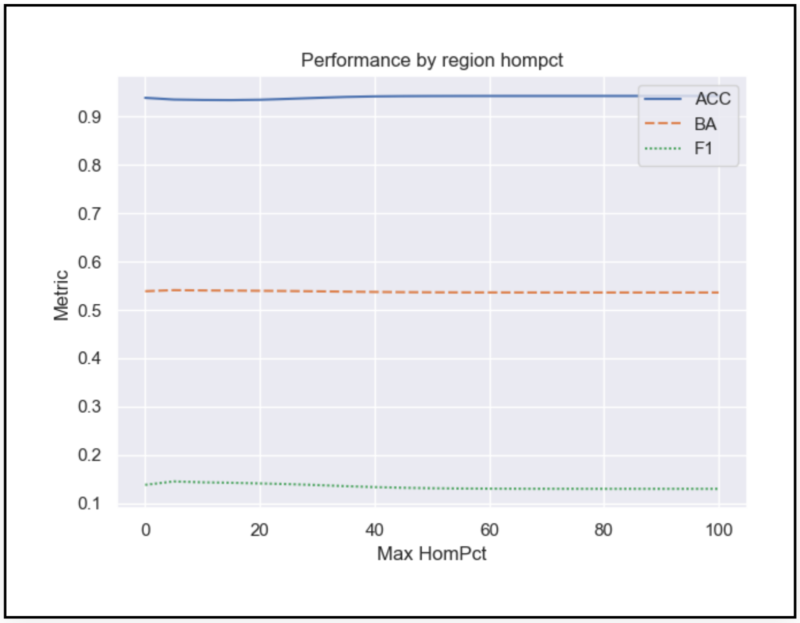
Performance by region hompct
ACC
BA
F1
Metric
Max HomPct

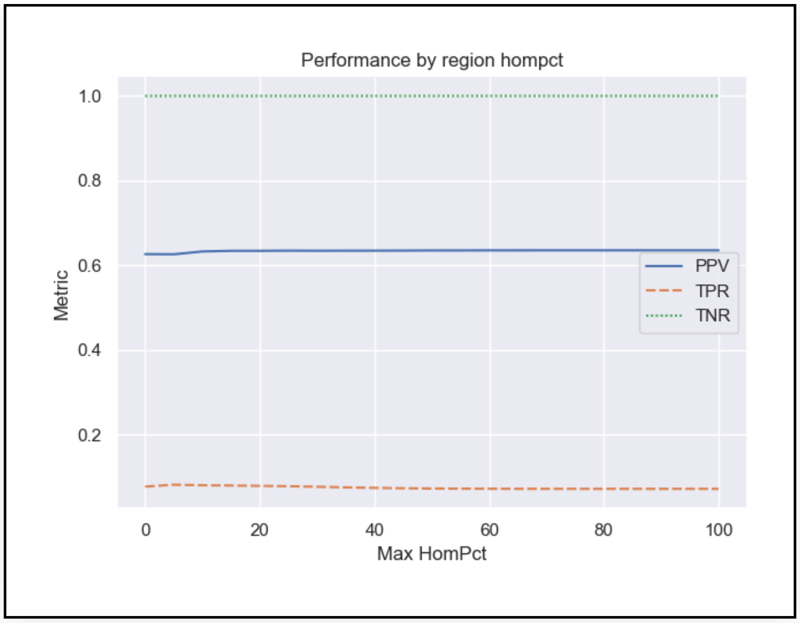
Performance by region hompct
PPV
TPR
TNR
Metric
Max HomPct

#### SOMs ?

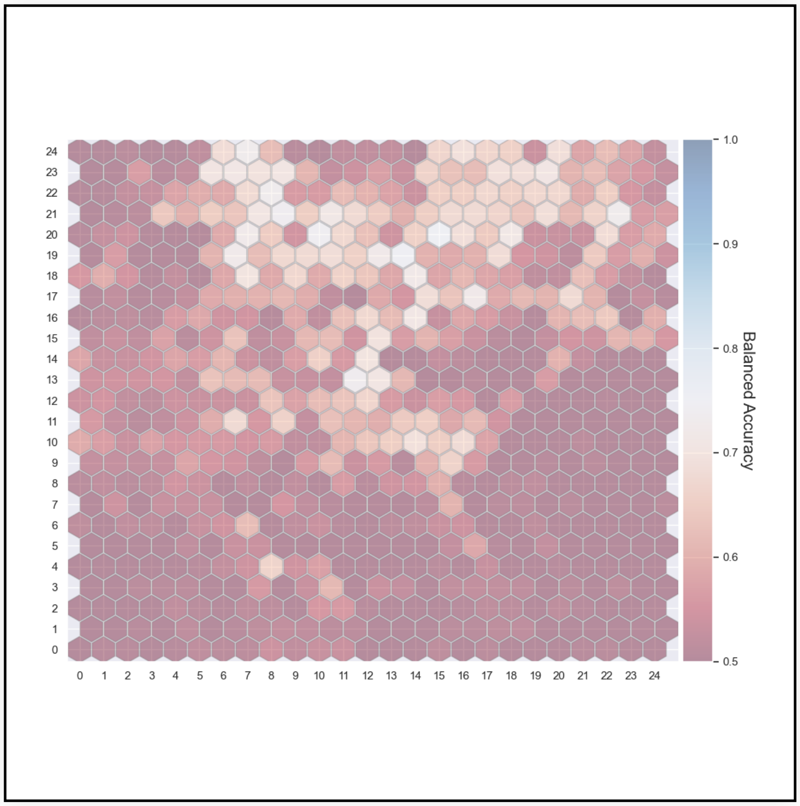
Balanced Accuracy

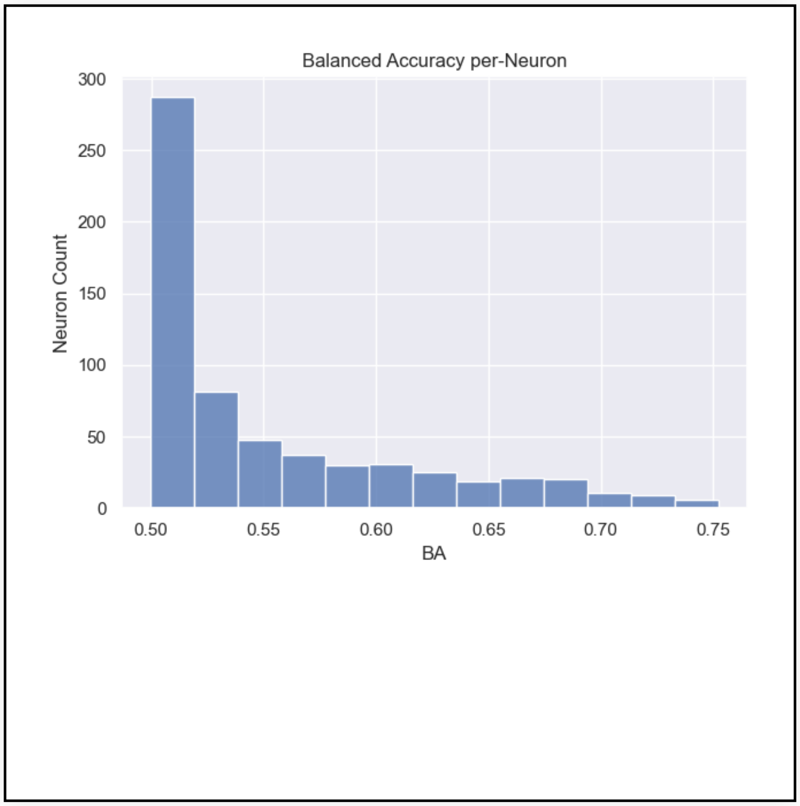
Balanced Accuracy per-Neuron
Neuron Count
BA

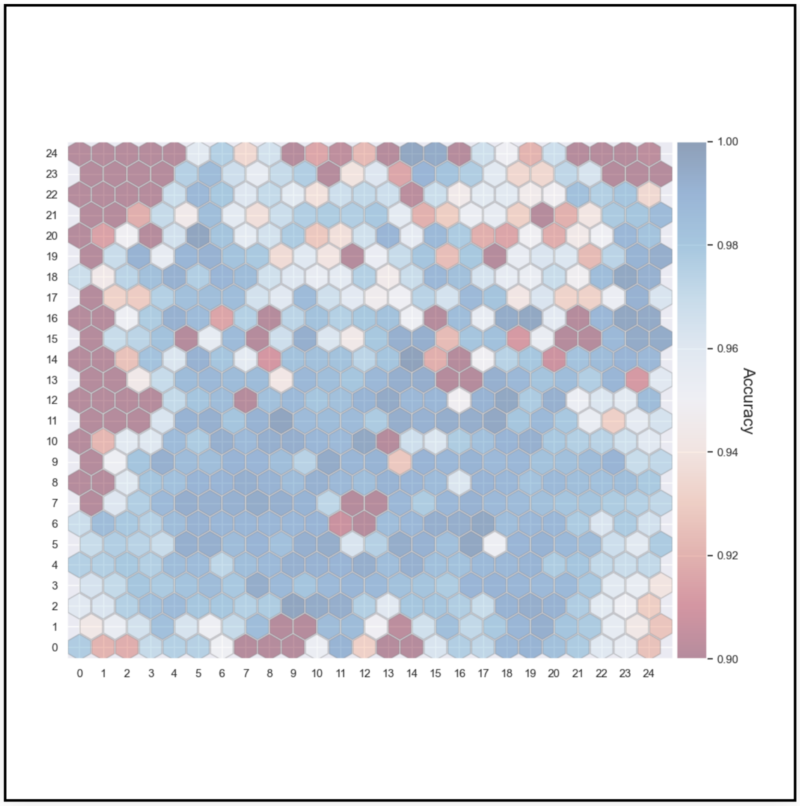


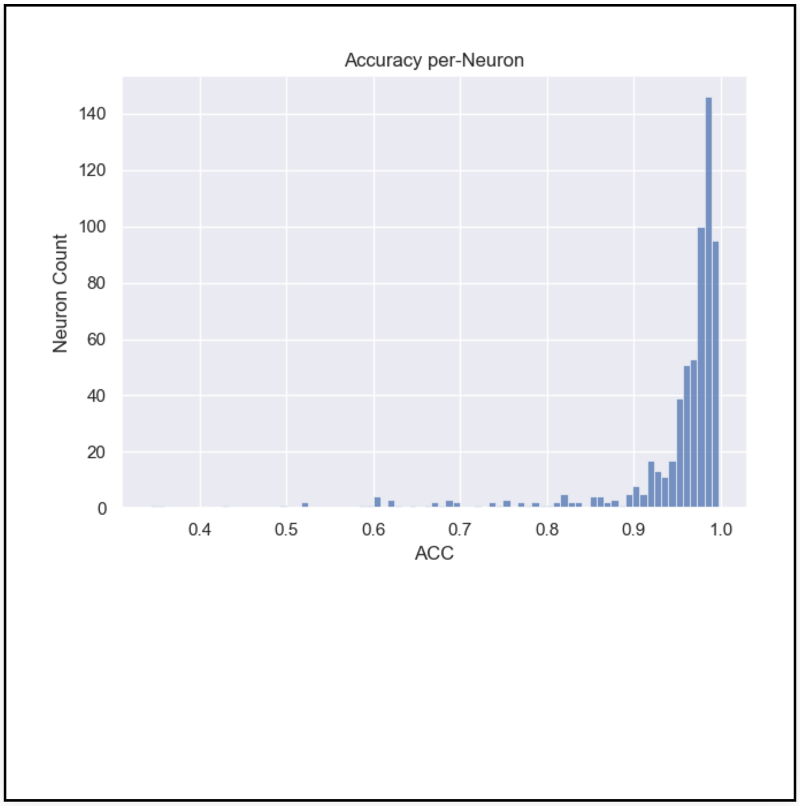


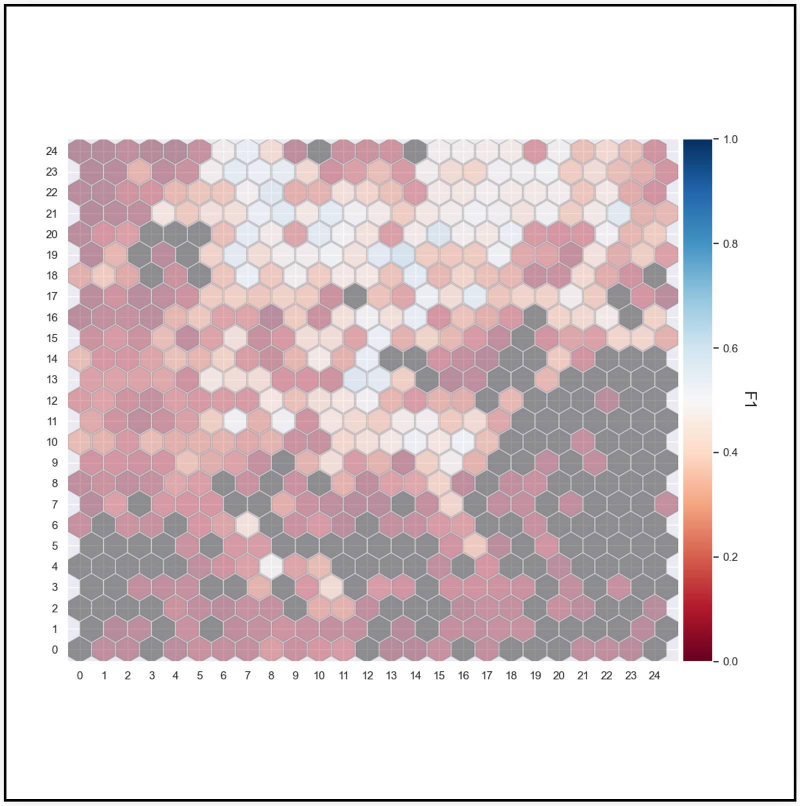


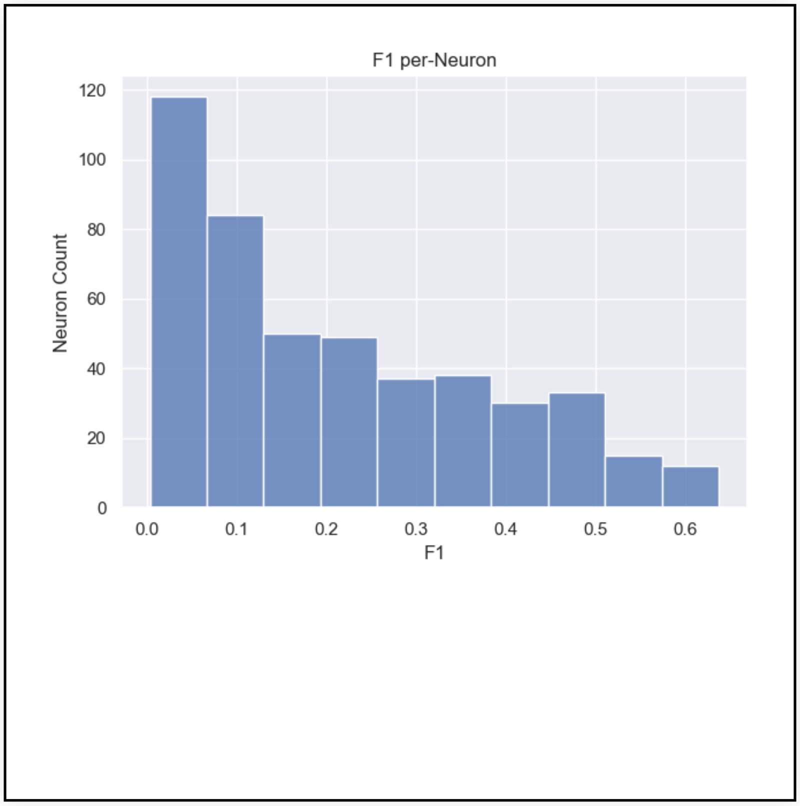


#### Reference Expansion Contraction Mixed* ?

Download

| | TP | TN | FP | FN | base P | base N | comp P | comp N | PPV | TPR | TNR | NPV | ACC | BA | F1 |
|---|---|---|---|---|---|---|---|---|---|---|---|---|---|---|---|
| **ExpCon** | | | | | | | | | | | | | | | |
| **EXP** | 7576 | 198933 | 666 | 97352 | 104992 | 198959 | 11860 | 292091 | 0.638786 | 0.072158 | 0.999869 | 0.681065 | 0.679415 | 0.536014 | 0.129668 |
| **REF** | 17 | 1402748 | 65 | 72 | 89 | 1402813 | 89 | 1402813 | 0.191011 | 0.191011 | 0.999954 | 0.999954 | 0.999902 | 0.595482 | 0.191011 |

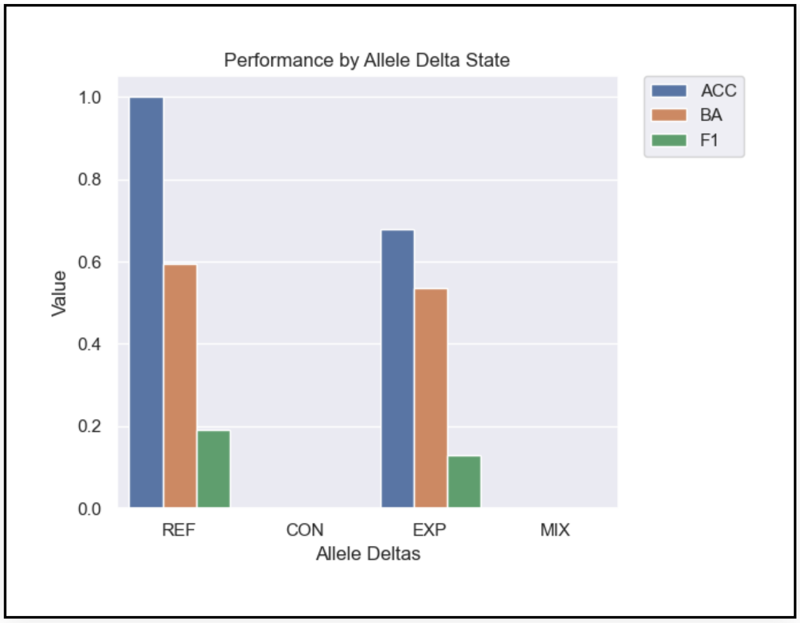


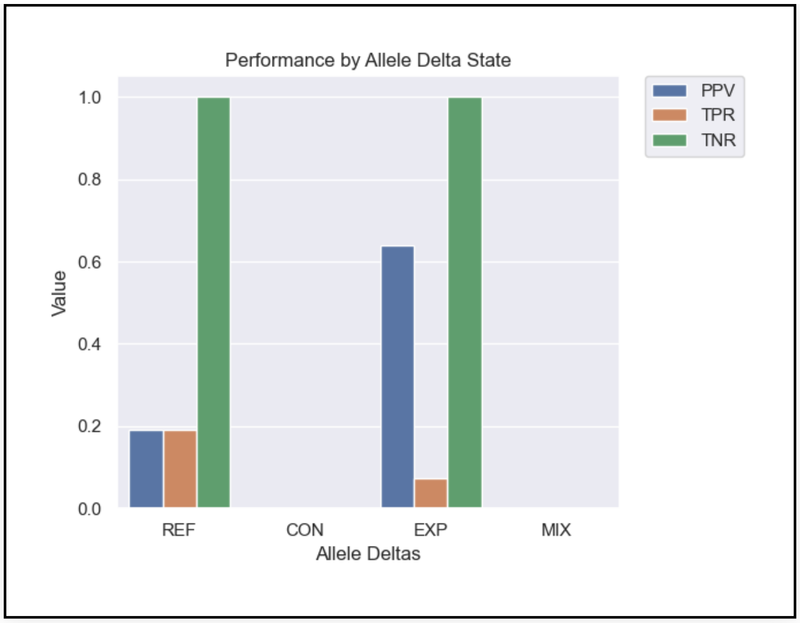


#### Max SizeBin* ?

Download

| | TP | TN | FP | FN | base P | base N | comp P | comp N | PPV | TPR | TNR | NPV | ACC | BA | F1 |
|---|---|---|---|---|---|---|---|---|---|---|---|---|---|---|---|
| **SNP** | 17 | 1402748 | 65 | 72 | 89 | 1402813 | 89 | 1402813 | 0.191011 | 0.191011 | 0.999954 | 0.999954 | 0.999902 | 0.595482 | 0.191011 |
| **[1,5)** | 10 | 192482 | 19 | 23 | 33 | 192498 | 33 | 192498 | 0.303030 | 0.303030 | 0.999917 | 0.999917 | 0.999797 | 0.651474 | 0.303030 |
| **[5,10)** | 5 | 4706 | 7 | 40426 | 40432 | 4709 | 20 | 45121 | 0.250000 | 0.000124 | 0.999363 | 0.104297 | 0.104362 | 0.499743 | 0.000247 |
| **[10,15)** | 1 | 794 | 4 | 22062 | 22063 | 795 | 10 | 22848 | 0.100000 | 0.000045 | 0.998742 | 0.034751 | 0.034780 | 0.499394 | 0.000091 |
| **[15,20)** | 1 | 325 | 2 | 10436 | 10437 | 325 | 6 | 10756 | 0.166667 | 0.000096 | 1.000000 | 0.030216 | 0.030292 | 0.500048 | 0.000192 |
| **[20,30)** | 3 | 371 | 6 | 11447 | 11451 | 371 | 21 | 11801 | 0.142857 | 0.000262 | 1.000000 | 0.031438 | 0.031636 | 0.500131 | 0.000523 |
| **[30,40)** | 1 | 140 | 7 | 5066 | 5067 | 140 | 20 | 5187 | 0.050000 | 0.000197 | 1.000000 | 0.026991 | 0.027079 | 0.500099 | 0.000393 |
| **[40,50)** | 25 | 62 | 3 | 2614 | 2641 | 62 | 62 | 2641 | 0.403226 | 0.009466 | 1.000000 | 0.023476 | 0.032186 | 0.504733 | 0.018498 |
| **[50,100)** | 2835 | 32 | 34 | 2123 | 4961 | 33 | 4017 | 977 | 0.705751 | 0.571457 | 0.969697 | 0.032753 | 0.574089 | 0.770577 | 0.631544 |
| **[100,200)** | 1920 | 0 | 75 | 1147 | 3073 | 0 | 2974 | 99 | 0.645595 | 0.624797 | NaN | 0.000000 | 0.624797 | NaN | 0.635026 |
| **[200,300)** | 747 | 1 | 57 | 452 | 1206 | 1 | 1192 | 15 | 0.626678 | 0.619403 | 1.000000 | 0.066667 | 0.619718 | 0.809701 | 0.623019 |
| **[300,400)** | 574 | 2 | 35 | 280 | 859 | 2 | 842 | 19 | 0.681710 | 0.668219 | 1.000000 | 0.105263 | 0.668990 | 0.834109 | 0.674897 |
| **[400,600)** | 505 | 7 | 52 | 326 | 838 | 10 | 820 | 28 | 0.615854 | 0.602625 | 0.700000 | 0.250000 | 0.603774 | 0.651313 | 0.609168 |
| **[600,800)** | 319 | 3 | 44 | 192 | 513 | 3 | 499 | 17 | 0.639279 | 0.621832 | 1.000000 | 0.176471 | 0.624031 | 0.810916 | 0.630435 |
| **[800,1k)** | 159 | 2 | 39 | 135 | 301 | 2 | 289 | 14 | 0.550173 | 0.528239 | 1.000000 | 0.142857 | 0.531353 | 0.764120 | 0.538983 |
| **[1k,2.5k)** | 369 | 6 | 165 | 410 | 790 | 8 | 759 | 39 | 0.486166 | 0.467089 | 0.750000 | 0.153846 | 0.469925 | 0.608544 | 0.476436 |
| **[2.5k,5k)** | 81 | 0 | 69 | 127 | 215 | 0 | 205 | 10 | 0.395122 | 0.376744 | NaN | 0.000000 | 0.376744 | NaN | 0.385714 |
| **>=5k** | 21 | 0 | 48 | 86 | 112 | 0 | 91 | 21 | 0.230769 | 0.187500 | NaN | 0.000000 | 0.187500 | NaN | 0.206897 |

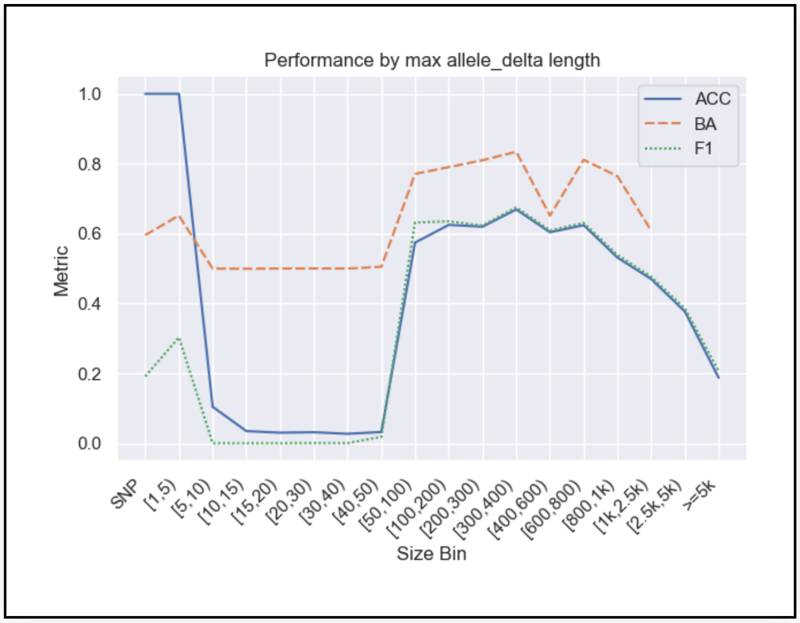


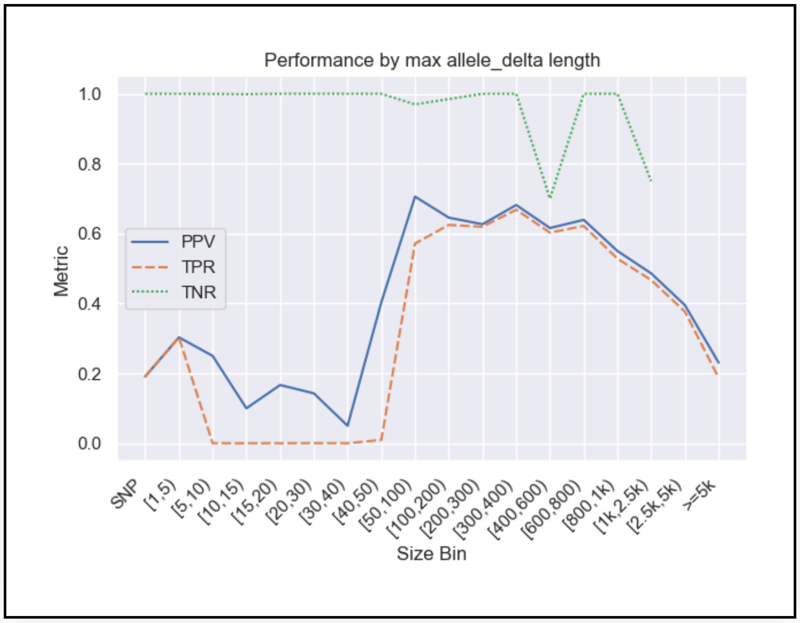


*Stratification by sizes requires a present variant. Therefore metrics which use TN (e.g. TNR, NPV, ACC) aren't informative.

---
